# Biological mechanisms are necessary to improve projections of species range shifts

**DOI:** 10.1101/2024.05.06.592679

**Authors:** Victor Van der Meersch, Edward Armstrong, Florent Mouillot, Anne Duputié, Hendrik Davi, Frédérik Saltré, Isabelle Chuine

## Abstract

The recent acceleration of global climate warming has created an urgent need for reliable projections of species distributions, widely used by natural resource managers. Such projections, however, are produced using various modeling approaches with little information on their relative performances under expected novel climatic conditions. Here, we hindcast the range shifts of five forest tree species across Europe over the last 12,000 years to compare the performance of three different types of species distribution models and determine the source of their robustness. We show that the performance of correlative models (CSDMs) decreases twice as fast as that of process-based models (PBMs) when climatic dissimilarity rises, and that PBM projections are likely to be more reliable than those made with CSDMs, at least until 2060 under scenario SSP245. These results demonstrate for the first time the well-established albeit so far untested idea that explicit description of mechanisms confers models robustness, and highlight a new avenue to improve model projections in the future.

## Introduction

Modelling is essential for understanding and predicting the impacts of climate change on ecosystems and biogeochemical cycles. Credible model projections are critical for natural resource managers, decision makers and stakeholders to make informed decisions. To meet the demand for reliable projections of ecosystems and biodiversity dynamics, comprehensive assessments of ecological model performances must be a priority (1–3).

One approach to evaluate model reliability is to compare their predictions to observations from previous time periods, i.e. hindcasting. Hindcasting can inform whether models capture, implicitly or explicitly, the essential processes required to provide reliable projections in conditions significantly different from the present. By looking far into the past, paleoarchives have proven to offer a unique opportunity to both understand long-term climate and biodiversity dynamics (4, 5) and test model robustness and transferability (i.e. model capacity to maintain its performance in changing conditions; 6) (7, 8).

Yet, models prediction of past species distribution and biospheric components have rarely been consistent with paleoclimate reconstructions and fossil records (9–13). Interpreting model projections in climatic conditions that differ significantly from the present, such as future no-analogue climatic conditions (14), remains challenging. Therefore, the guarantee that ecological model forecasts for the 21^st^ century will be reliable is limited (15).

While exact matches to expected 21^st^-century climatic conditions do not exist in historical records (16), the dissimilarity between 20^th^ and 21^st^ century median climatic conditions (Methods) falls within the range of dissimilarity encountered since the beginning of the Holocene (12 kyr BP, with kyr = 1000 years and BP = Before Present (1950); Figure 1). This period takes place after the Last Glacial Maximum (26.5-19 kyr BP; 17) and began with an abrupt climate warming followed by a long, almost uninterrupted, period of climatic stability until recent anthropogenic warming (Figure S1). The fossil pollen data accumulated over these last millennia provides us with an unique extended timeframe to test the reliability of ecological models, in particular those designed to predict changes in species distribution.

**Fig. 1.**
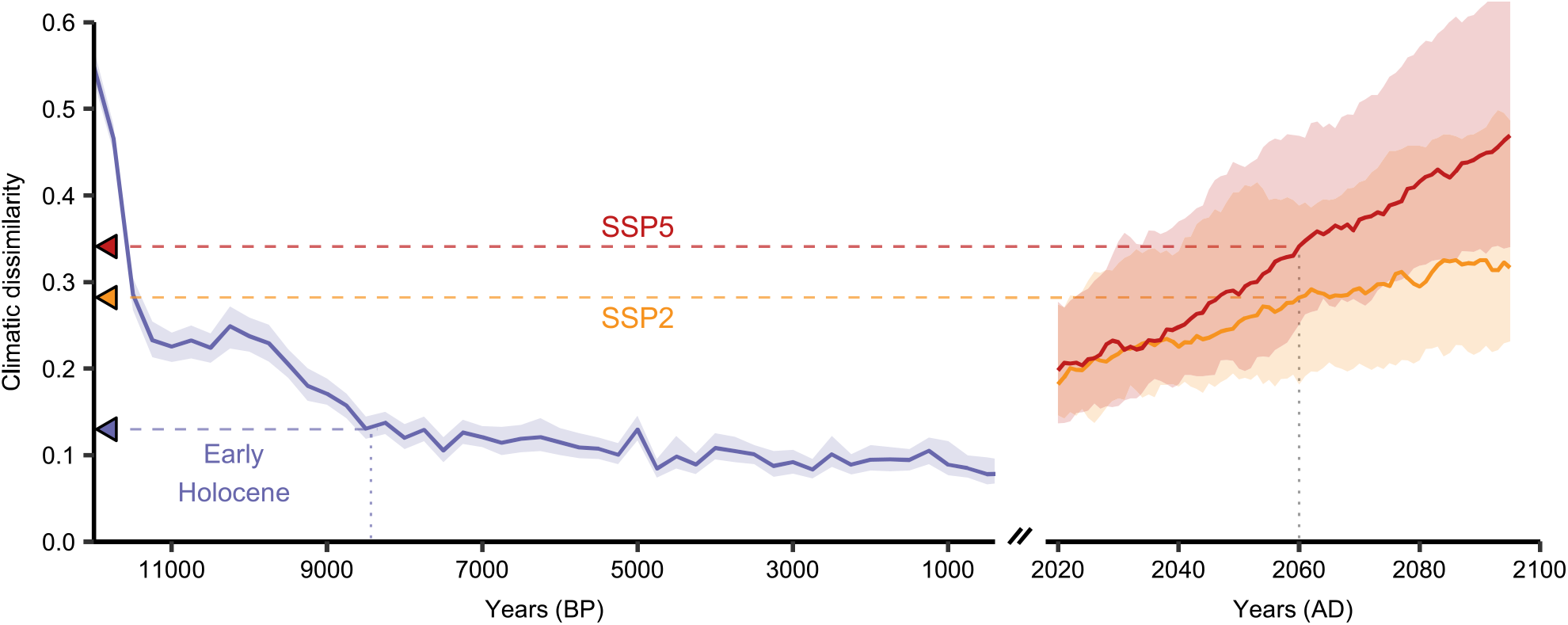
Evolution of climatic dissimilarity during the Holocene (12k-500 yr BP) and the 21st century (2005-2100), relative to 1901-2000. Climatic dissimilarity is computed as 1-Sørensen similarity between bootstrapped climatic hypervolumes. Lines represent median dissimilarity, shaded areas represent 90% confidence intervals. Blue corresponds to paleoclimate based on HadCM3B model. Yellow and red correspond to future climatic conditions according to SSP245 and SSP585 scenarios respectively, predicted by 34 global climate models of NEX-GDDP-CMIP6. The blue triangle on y-axis indicates the level of climatic dissimilarity 8500 years ago, at the limit between the Early and Late-Middle Holocene. Yellow and red triangles indicate the expected level of climatic dissimilarity in 2060 for SSP245 and SSP585 scenarios. Note that the x-axis scale is different between past and future panels.

Species distribution models (SDM) are powerful tools to predict species geographical distribution as a function of environmental data (e.g., mean annual temperature and annual total precipitation). Most studies have focused on correlative models (CSDMs, also called environmental niche models), which infer statistical relationships between observations of species occurrences and environmental predictors (18). Their high flexibility and low computational complexity make them the most widely used tool for deciding on species conservation plans and policy regimes (e.g. 19). However, under novel climatic conditions, new unobserved portions of a species’ climatic niche may appear, which are not captured by these correlative (or phenomenological) approaches. For example, when tested under distant past climates, the predictive performance of CSDMs significantly decreased (13), questioning their ability to provide reliable projections in the future (15). Alternative approaches to CSDMs are process-based models (PBMs, or process-explicit models) that rely on explicit formulations of the mechanisms driving the distribution of a given species (e.g., physiological, ecological and/or demographic processes). They come from decades of experiments and observations, including extreme conditions in laboratory (20), and climate manipulations such as CO_2_ enrichment (21) or rainfall exclusion (22). The reliability of PBMs depends on our level of understanding of how environmental conditions affect ecophysiological processes, and the availability of large amount of observations to calibrate their many parameters (23). Because these models do not rely on statistical relationships between present-day species occurrences (presence/absence) and environmental variables, but rather describe explicit causal relationships between biological processes and environmental variables, they are believed to provide more reliable predictions of species distribution changes under novel climatic conditions (24, 25). However, another possible reasons why PBM projections might be more reliable than CSDM projections under novel climatic conditions could also come from their calibration methods. Unlike CDSMs that are calibrated using species presence/absence data, PBM parameters are either measured directly (e.g. specific leaf area, leaf frost hardiness), or inferred statistically when direct measurement is not an option, using data on specific functional traits measured in the field or in laboratory (e.g. parameters of bud dormancy break date models).

The assumption that PBMs could provide more reliable projections of future range shifts of species is widely accepted and taken for granted (24, 26–28) although it has never really been demonstrated. Furthermore, the reasons behind this assumption have not been clearly articulated. Qualitative models comparison under future climatic conditions have shown that PBMs often make more conservative projections in future climates than CSDMs which predict larger changes (29–31) but they have not provided any confidence level in these results. Very few studies have actually gone beyond qualitative comparisons between CSDMs and PBMs and compared thoroughly their performance, for example using virtual species (32), exotic species in native and newly colonized areas (33), or in the recent past (34). While PBMs have shown their usefulness for paleoecological studies (35–37), the extent to which they can provide more reliable predictions than CSDMs under different climatic conditions from the historical period remains unknown (6, 38).

Here, we address this critical gap by using multiple CSDMs and PBMs to simulate paleodistributions of five emblematic tree species of Europe at a high temporal resolution since 12 kyr BP. We used daily paleoclimatic data at 0.25° spatial resolution, generated from HadCM3B-M2.1 coupled general circulation model simulations, which includes both inter-annual variability, and millennial scale variability for rapid Dansgaard–Oeschger events before 11 kyr BP (39). Species migration ability was also incorporated into the simulations to represent more comprehensively changes in species’ realized distribution and not merely changes in their climatic niches to allow for a more accurate comparison with the paleorecords. We first assessed which modelling approach best predicts past species distributions, and second whether model performance was related to their hypotheses (relationships describing explicit biological mechanisms or not) or to their calibration methods (calibrated on species occurrence data or not). To do so, we compared three types of models: CSDMs, PBMs (hereafter called expert PBMs) and fitted PBMs calibrated in the same way as CSDMs (inverse calibration using species occurrence data and a novel type of algorithm; Methods; 40). The comparison between CSDMs/fitted PBMs and expert PBMs allowed us to determine whether the differences in model performance arise from their calibration methods, whereas the comparison between CSDMs and expert/fitted PBMs allowed us to determine whether the differences in model performance arise from the model hypotheses. We assessed the performance of the models for the maximum level of climate dissimilarity possible, i.e. over the last 12,000 years, which corresponds to the climate dissimilarity expected by the end of the century according to SSP245, and by the middle of the century according to SSP585 (Figure 1).

## Methods

### Correlative and process-based species distribution models

We used PHENOFIT and CASTANEA, two process-based models which differ by their underlying hypothesis and complexity. PHENOFIT simulates the fitness of an average adult tree (41). It estimates fitness components (survival and reproductive success) by simulating the precise phenology (dates of leaf unfolding, flowering, fruit maturation, leaf senescence) and damages caused by abiotic stress (frost, drought) which effects depend on their occurrence relatively to the development stages of the different organs. It has been validated for several North American and European species (35, 42–44). The model has ∼30 parameters. CASTANEA simulates carbon and water cycles of an average adult tree by simulating many processes such as photosynthesis, stomatal opening, maintenance and growth respiration, transpiration and carbon allocation (45). It has been used to predict carbon and water budgets of several European species (46–48). The model has ∼80 parameters. Both models require daily meteorological variables and soil characteristics. Two versions of both models were employed: one was calibrated with expert knowledge and statistical inference using observations and measurements of the processes modelled (version called *expert*), and a second one was entirely calibrated using species distribution data (version called *fitted*; 40) like correlative models. For the latter, we used the optimization algorithm CMA-ES (49) as described in (40), and retained the best calibrations in terms of AUC.

We selected correlative models based on the thorough model comparison made by (50). Among the most performant models, we selected five well-established models: GLM with lasso regularization, GAM, BRT, MaxEnt and down-sampled Random Forest. Some of these models are known to struggle when applied to extrapolation domains, but are nevertheless widely used by ecologists to provide projections of species distribution change in future climatic conditions. We chose four uncorrelated climate predictors related to ecological processes to calibrate these models: minimum temperature of the coldest month (representing frost tolerance), total precipitation (representing available water), GDD5 (growing degree days >5°C) between April and September (representing available thermal energy for vegetation growth and fruit maturation), water balance between June and July (precipitation*−*evapotranspiration, representing summer drought tolerance). We also included two soil covariates (pH and Water Holding Capacity).

While by construction correlative models directly output species habitat suitability, we used fitness predicted by the model PHENOFIT and net primary production predicted by the model CASTANEA as a proxy of species habitat suitability as they have already been used to predict species presence in previous studies (29, 30, 35). CSDMs and inverse-calibrated PBMs were calibrated for five species (*Fagus sylvatica* L., *Abies alba* Mill., *Quercus robur* L., *Quercus petraea* (Matt.) Liebl. and *Quercus ilex* L.) using historical climate (1970-2000) extracted from ERA5-Land hourly dataset (51), soil data from EU-SoilHydroGrids (52) and SoilGrids (53) databases and species presence data from the dataset assembled in (40), mostly based on EU-Forest inventory data (54). To calibrate the CSDMs, we additionally sampled 50,000 background points, which should properly represent the variation in the environmental conditions across the study area (50). For each CSDM and each species, we run a five-fold environmental cross-validation to estimate model performance in novel extrapolation conditions (Figure S8; 55). We then used all the available training data to calibrate the models for the hindcasting in order to favour final prediction quality (55). We could not run the same cross-validation method for fitted process-based models because it would have been too computationally expensive.

Model simulations over the Holocene were run for 30-year periods every 250 years, for the five above mentioned species. Model outputs were averaged over each 30-year period. Note that soil conditions (needed both for correlative and process-based models) were held constant throughout the simulations, and were bilinearly interpolated from closest coastal cells where data was missing (because of different land-sea masks between present and past). Note also that for CASTANEA model, species-specific thresholds of net primary production determining the presence or absence of the species were computed with the CO_2_ level at the beginning of the Holocene (∼240ppm).

### Late Quaternary climate and vegetation

We used the monthly paleoclimate simulation dataset generated with the HadCM3B-M2.1 coupled general circulation model (39), starting from 18 kyr BP at 0.5° spatial resolution for Europe (Figure S1). We chose this dataset for several reasons. First, it includes both inter-annual variability, and millennial scale variability for rapid Dansgaard–Oeschger events before 11 kyr BP. Second, it shows generally a good agreement with ice-core datasets (39). Third, it provides all the necessary input variables necessary to run all the models selected. For this work, several variables were specifically produced: mean temperature, average minimum and maximum daily temperatures, precipitation, number of rainy days, cloudiness, and wind speed. We further downscaled temperature and precipitation monthly data to 0.25° resolution, by applying an elevation correction of coarse-scale variables towards the ICE-6G-C elevation level at high resolution (56). We then generated daily data for temperatures, precipitation, cloud cover and wind speed from the monthly data with the weather generator GWGEN (57), for 30-year periods every 250 years. We also simulated daily extra-terrestrial solar radiation with the same orbital forcing conditions used in HadCM3B-M2.1 (39) and then computed daily global radiation taking into account previously generated daily cloud-cover data as implemented in LPJ-LMfire global model (58). Finally, we computed daily potential evapotranspiration following the standard FAO Penman-Monteith method (59).

Fossil pollen records were extracted from the Legacy-Pollen dataset (60). This dataset is mainly based on the Neotoma database (61), and provides samples with standardized chronologies and age uncertainties. We removed sites that had marine depositional environments (13), and only kept samples with more than 200 pollen grain counts and age uncertainty of less than 500 years. Pollen relative abundances were aggregated to consecutive 500-year intervals. If multiple samples from the same site belonged to the same period, we averaged their pollen abundances, weighting by their age uncertainty and temporal distance from the center of the period. Relative pollen abundances were converted to presence/absence using thresholds based on biome reconstructions (62): 1% for *Fagus* and *Abies*, and 2.5% for *Quercus*. If several sites fell within the same grid cell (0.25°), we considered the species as present if there was at least one site where the species could be considered as present. *Fagus* pollen data were used to assess the presence of *Fagus sylvativa* L., sole species of the genus present in Europe. *Abies* pollen data were used to assess the presence of *Abies alba* Mill., most abundant and widespread fir species present in Europe. When possible, deciduous and evergreen *Quercus* pollen were distinguished based on Neotoma data. Some *Quercus* pollen remain undetermined beyond the generic level, either because discrimination between evergreen and deciduous oak pollen was impossible or because authors were not specific. They were assigned to two categories, based on the evergreen natural range as defined by Atlas Flora Europeae (63) and EuroVegMap (64): pollen outside range were considered as deciduous only occurrences, whereas pollen inside range were considered as both evergreen and deciduous occurrences. Deciduous *Quercus* pollen data were used to assess the presence of *Quercus petraea* (Matt.) Liebl., and *Quercus robur* L., the two most abundant and widespread deciduous oak species in Europe. Evergreen *Quercus* pollen data were used to assess the presence of *Quercus ilex* L., the most abundant and widespread evergreen oak species in Europe.

### Tree migration

Models used in this study predict species potential distribution based solely on climatic and soil conditions. To compare model predictions to pollen paleorecords, species migration needs to be simulated as well, as it can be the primary factor limiting species distribution before climatic conditions, especially when climatic conditions are changing rapidly as it was the case during the Dansgaard–Oeschger events (35, 65).

To implement migration in the simulations, we run a cellular automaton (66) which has proven to be as accurate as more complex approaches (32). We modified the initial version of this dispersal model in order to use both short- and long-distance dispersal kernels (long distance events could occur with a probability of 0.01). We used species-specific fat-tailed kernels (67) at a 500 m resolution, and assumed that trees can disperse once a year (Figure S7a). Model outputs were assigned to two classes using specific optimal thresholds maximizing model performance (TSS) in the historical climate (Figure S8): (i) cells where the model output was under the specific threshold were assigned a zero suitability (species cannot survive), and (ii) cells where the model output was above the threshold, the suitability was rescaled between 0 and 1 (species can migrate). We considered the deciduous *Quercus* suitability as the maximum suitability between *Q. robur* and *Q. petraea*. Migration simulations started from 12 kyr BP (or 11.75 kyr BP when a model simulates no presence at 12 kyr BP). Note that starting at 11.75 kyr BP or 12 kyr BP does not change our results (Figure S10e-h), and that we could not start earlier (e.g. ∼15 kyr BP) as most models predict no presence at all around 12.5 kyr BP. We also checked that dispersal process stochasticity at 500m resolution (Figure S7a) had no significant effect on the model’s performance at the scale of Europe, by simulating deciduous *Quercus* migration 10 times for each of the 8 models (Figure S7b).

### Models’ performance

We used the Sørensen’s similarity index to measure the hindcast performance of the models, based on the confusion matrix. This discrimination measure has been shown to provide adequate estimations of model discrimination capacity, not biased by species prevalence or an inflated number of true negative predictions (68). This feature is important when working with fossil pollen data, for which the number of species absence can be much higher than the number of species presence. Note that we obtained similar results when using TSS as the performance metric (Figure S10b). We compared the area potentially occupied (not taking migration into account) and occupied (taking migration into account) by the species to the presence/absence data extracted from the LegacyPollen dataset every 500-year interval. Kruskal-Wallis tests followed by multiple pairwise post-hoc Conover-Iman tests (using Benjamini-Yekutieli adjustment, as implemented in the R package *conover*.*test*) were computed to assess stochastic dominance among model performance and transferability (Fig. 3b-d).

To quantify the climatic differences between historical climate (1901-2000, based on the CRU TS v. 4.07 gridded dataset; 69) and Holocene climate (hindcasting conditions), we computed the *climatic dissimilarity* as the Sørensen dissimilarity between climatic hypervolumes (a metric of overlap in multidimensional space). We first generated for each period (500-year intervals from -12 kyr BP to 500 BP and 1901-2000) a set of 20 bootstrapped hypervolumes, using R package *hypervolume* (70). Hypervolumes were computed with a Gaussian kernel density estimation method based upon the first three principal component axis from three-month means temperature and three-month sums of precipitation. We then computed overlap statistics (mean and standard deviation of Sørensen index) between the bootstrapped hyper-volumes of each time points of the Holocene and the boot-strapped hypervolumes of the historical period (i.e. 20x20 overlaps). As a comparison, we also computed the climate novelty based on Mahalanobis distance (Figure S2; 71).

We also computed these metrics under future conditions to compare the dissimilarity of future climate to that of the Holocene climate, both relative to 20th century climate. To assess future conditions, we used all the global climate models from NEX-GDDP-CMIP6 dataset (72) – except HadGEM3-GC31-MM, not available for SSP245 – and 2 scenarios (SSP245 and SSP585). To make the comparison, both paleoclimate and future climate data were uniformized with the CRU dataset to maximize comparability between paleo-climate and future climates. The difference (for three-month temperature average) and the ratio (for three-month precipitation sum) between the observations (from 1901 to 2000) and simulations (1901-1950 for HadCM3B, 1951-2000 for CMIP6 projections) were calculated and applied to the whole modelled time period, assuming that the bias was constant. Finally, we estimated the effects of past climate novelty (Sørensen’s climatic dissimilarity) on model performance (Sørensen index) with a Bayesian ordered beta regression, considering the different types of models (correlative, fitted process-based and expert process-based), using the R package *ordbetareg* (73) and RStan (74). Compared to a standard beta regression model, this model allows for observations at the bounds (i.e. Sørensen index = 0 or = 1). We took into account the standard deviation of Sørensen’s climatic dissimilarity (computed with sets of bootstrapped hypervolumes, see above) as a predictor measurement error.

## Results

As observed in previous long-term historical assessments, all models showed a decrease of their performance when moving further into the past, i.e. into more different climatic conditions than historical conditions (Fig. 3a). However PBMs showed smaller decrease in their predictive performance (slope of Beta regression, fitted PBMs: *−*6.07, 95% CI [*−*8.62, *−*3.55], expert PBMs: *−*4.44, 95% CI [*−*7.07, *−*1.77]) than CSDMs (*−*11.0, 95% CI [*−*13.2, *−*8.91]). PBMs also showed higher transferability in the most distant climatic conditions of the early Holocene than CSDMs (Fig. 3d). PBMs, either expert or fitted, are thus less affected by the increase in climate dissimilarity than CSDMs. In the near past (Late-Middle Holocene, *<* 8.2 kyr BP), CSDMs were not significantly better at predicting tree distribution than any PBMs (pairwise Conover-Iman tests: *vs*. expert PBMs *t*-statistic = 1.68/*P* = 0.128, *vs*. fitted PBMs *t*-statistic = *−*1.55/*P* = 0.112; Fig. 3b), despite their closer fit to current species distributions (Figure S8). In the distant past (Early Holocene *>* 8.2 kyr BP), CSDMs performed worse than both expert and fitted PBMs (pairwise Conover-Iman tests: respectively *t*-statistic = *−*4.80/*P <* 0.0001 and *t*-statistic = *−*5.07/*P <* 0.0001; Fig. 3b). The maximum climatic dissimilarity during this period corresponds to the climatic dissimilarity expected as soon as 2060 according to the scenario SSP245 (Fig. 3a).

These differences between PBMs and CSDMs are closely related to their ability to predict species recolonisation dynamics in the Early Holocene (∼11.5-8.5 kyr BP). Both types of PBMs, fitted and expert, predicted more accurately refugia locations at -12 kyr BP, which were the starting points for the migration (Fig. 2; Methods). This period corresponds to a global deglaciation which lasted for a few centuries and occurred after the cooling of the Younger Dryas interval (∼13-11.5 kyr BP; Figure S1). This rapid warming episode explains the strong decrease of climate dissimilarity relative to present between 12 kyr BP and 11.5 kyr BP (Fig. 1). If we had not considered the 12-11.5 kyr BP period of high climatic dissimilarity (i.e. without simulating migration from refugia), we would have missed the opportunity to take into account model projections within the same dissimilarity level to what we expect between 2050 and 2100 (Fig. 1). When models are compared after 11kyr BP, i.e. when climate dissimilarity is more similar to present, CSDMs and PBMs’ abilities to predict fossil pollen occurrence are similar (Figure S10cd), with an average Sørensen index decrease of *−*0.205 *±* 0.0224 (paired Wilcoxon-test: *P <* 0.0001) between Late-Middle Holocene and Early Holocene.

**Fig. 2.**
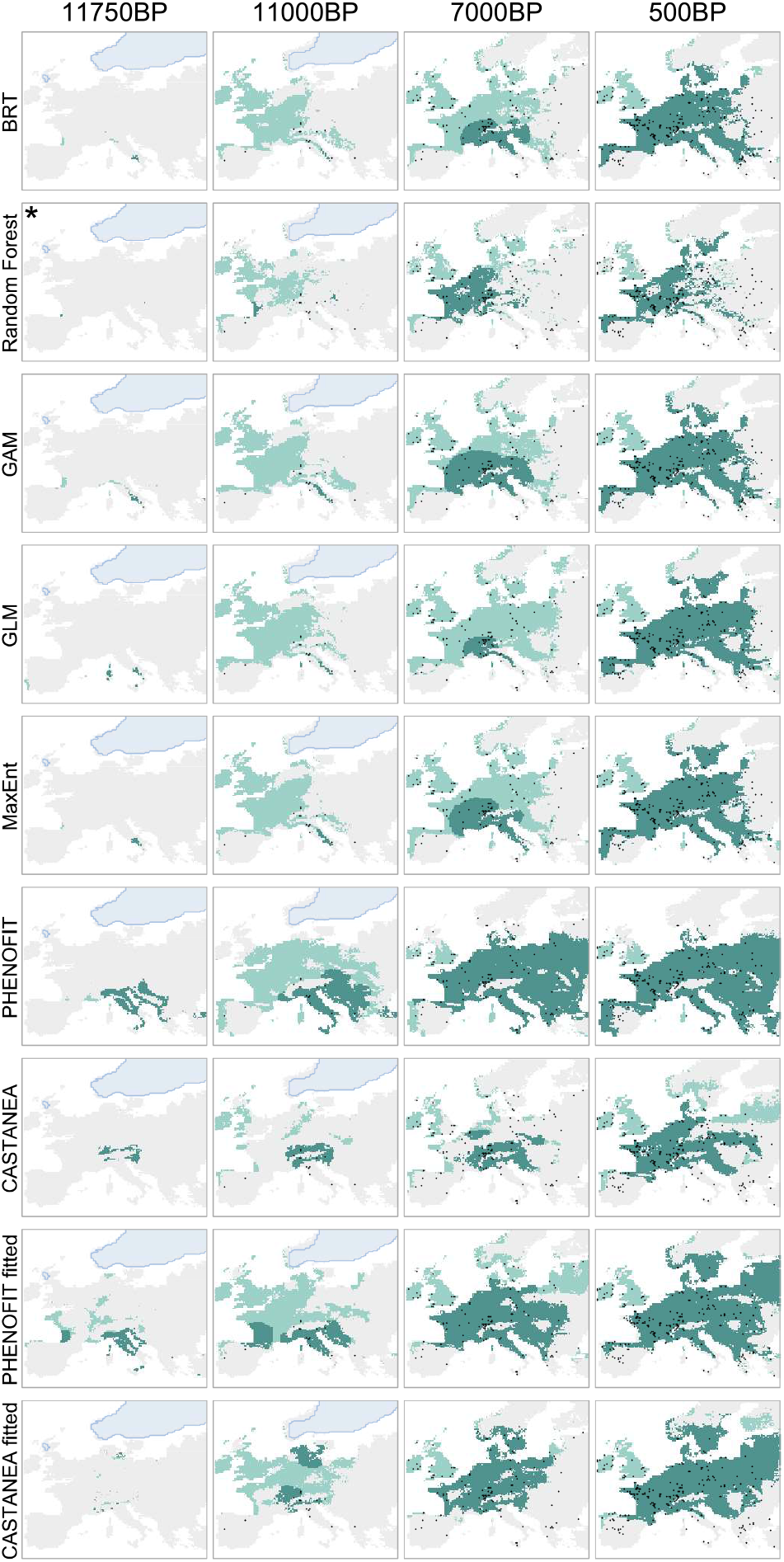
Example of paleosimulations obtained with the nine models used in this study for deciduous oaks. The five first rows correspond to the five correlative models (boosted regression tree, down-sampled random forest, generalized additive model, generalized linear model with lasso regularization, MaxEnt). The four last rows correspond to two different versions (expert calibration and inverse calibration using occurrence data) of two process-based models (PHENOFIT and CASTANEA). Light green area is the modelled suitable area, dark green area is the colonized area (after migration). Light blue represents the ice sheet extent. Black dots are deciduous oak fossil pollen occurrences. The model for which migration started at 11.75 kyr BP rather than 12 kyr BP is marked with an asterisk. “BP” stands for “before present” (1950).

Our results also revealed that inverse calibration improved process-based projections in recent past without altering significantly PBM long-term transferability (Fig. 3). In Middle-Late Holocene, when climate conditions were not drastically different from present, performances of fitted PBMs was higher than those of expert PBMs (*t*-statistic = 2.70/*P* = 0.020). In most distant climatic conditions of Early Holocene, their performances were similar (*t*-statistic = 0.220/*P* = 0.757; Fig. 3b).

**Fig. 3.**
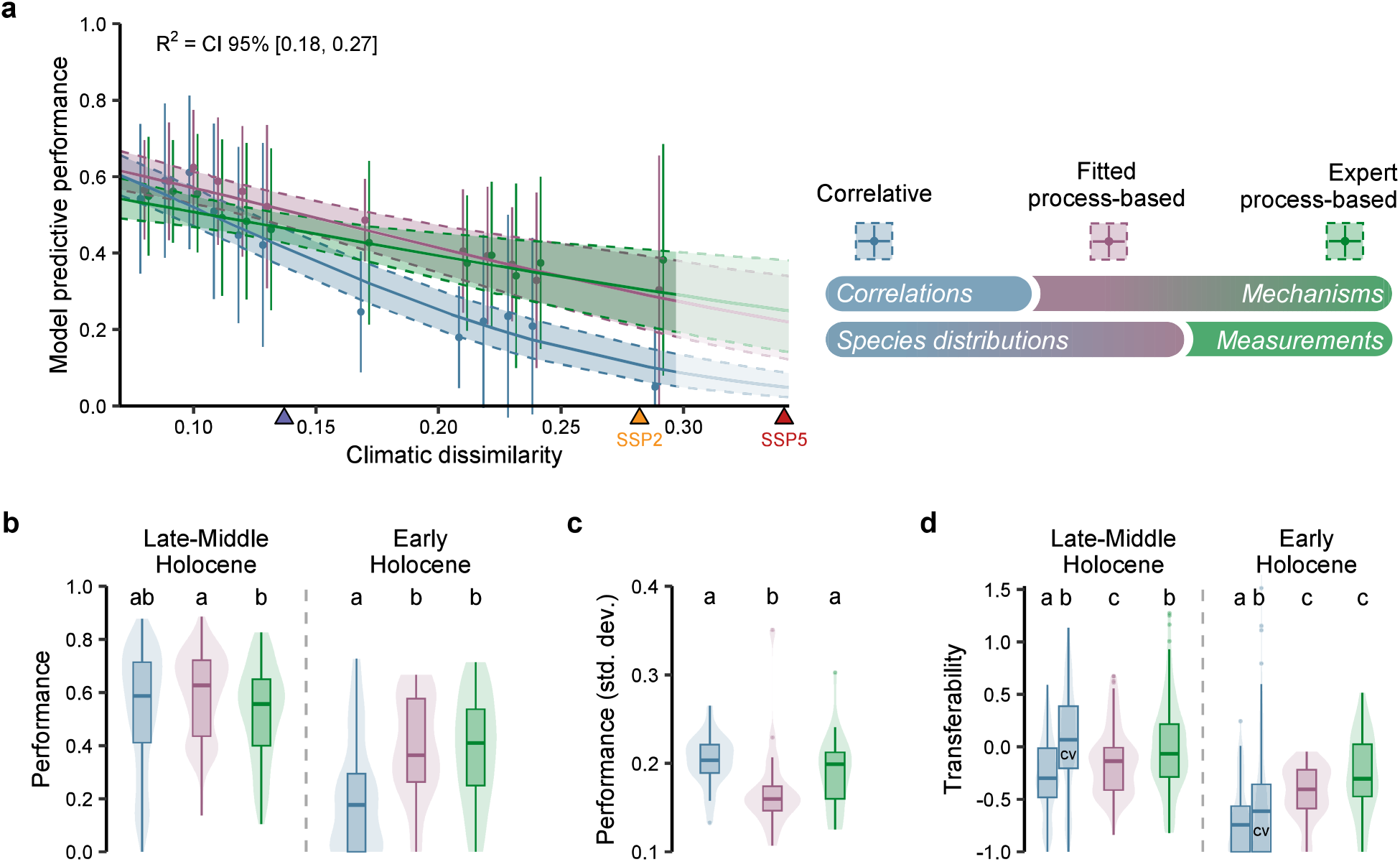
Performance of correlative models, fitted process-based models (inverse calibration using occurrence data) and expert process-based models (classical calibration) against Holocene paleoecogical evidence (fossil pollen) for 4 tree genera (*Abies, Fagus, Quercus* deciduous and *Quercus* evergreen). **(a)** Bayesian beta regression of model predictive performance (Sørensen index) against climatic dissimilarity relative to 1901-2000 (1-Sørensen similarity between climatic hypervolumes). Shaded areas represent 2.5% and 97.5% quantiles of the posterior predictive distribution. Points represent the average model performance (and lines the standard deviation) grouped by similar level of climatic dissimilarity. Blue triangle on x-axis indicates the limit between Early Holocene (*>* 8.2 kyr BP) and Late-Middle Holocene (*<* 8.2 kyr BP). Yellow and red triangles indicate the expected level of climatic dissimilarity in 2060 for SSP245 and SSP585 scenarios. Legend on the right: top row represents drivers of modelled distributions (correlations/mechanisms), bottom row represents calibration method (species distributions/measurements). Panels **(b)** and **(c)** show the difference in performance (Sørensen index) and variability in performance (standard deviation of Sørensen index) across models. Panel **(d)** shows the transferability of the models (relative change in model performance between distant past periods and historical period). A negative transferability means that model performance is lower in the distant past periods than in the historical period. CSDM predictive errors in the historical period was assessed by two different methods: (i) against the same data used for calibration (leading to an overestimation of historical model performance – but more comparable to fitted PBM estimates), (ii) using an environmental block cross-validation, noted as “CV” (leading to a better estimation of true model errors in the historical period, and thus a higher transferability – but less comparable to fitted PBMs for which cross-validation would have been too computationally expensive). The grouping letters represent the multiple comparisons with pairwise Conover-Iman tests.

Models performances were not stable across species, and exhibited both similarities and differences across time (Figure S9). More specifically, models exhibited the same overall performance decrease against *Fagus* pollen records, whereas CSDM performance decline was substantially faster than expert and fitted PBMs for deciduous *Quercus*. All models show low predictive power regarding evergreen *Quercus* distribution even in the Late Holocene compared to other species, especially CSDMs which failed to predict its presence along the Atlantic coast (S6). Fitted PBMs, however, showed the lowest variability of performance across species (Fig. 3c).

## Discussion

Our results suggest that the transferability and robustness of models are more strongly influenced by the processes explicitly represented in the models than by their method of calibration. PBMs consistently show a better performance throughout the 12 kyr simulation period, even when calibrated using the same method as CSDMs (i.e. fitted PBMs). Therefore, beyond enabling a more detailed mechanistic understanding of the effects of environmental conditions on species survival, growth, and reproduction, biological processes represented in PBMs are also critical to ensure higher model robustness under more novel climatic conditions. This important new finding advocates for a wider use of PBMs to predict biodiversity and ecosystems distributions in the future and opens a new avenue to reach this goal by using inverse modelling approaches to calibrate them.

Our results also suggest that predictions of PBMs, either fitted or expert, should be more reliable at least up to 2060 according to the scenario SSP245 (Fig. 3a) and 2050 according to SSP585. The rate of anthropogenic climate change and the increased probability of occurrence of novel climates (Fig. 1; 14) are challenging the reliability of both CSDMs and PBMs especially since they are intended to be used in more complex models such as biosphere-atmosphere models and used by natural resource managers and policy makers to guide management plans and policies. Acknowledging these uncertainties is as important as making the forecasts themselves (75) and contributes to the public trust in scientists (76).

Simulating migration allowed us to take into account the differences between the models under the most challenging conditions, i.e., when the climate dissimilarity was at its greatest, closely approximating what is projected for the end of the 21st century. Since the migration model is identical across all simulations, differences of performance between models across the Holocene very much depended on their ability to predict the potential refugia of the species during the Early Holocene. For example, some models were not able to predict evergreen *Quercus* refugia in Southern Spain, thus missed an important migration route and failed predicting their presence in vast areas in the Late Holocene (Figure S6). As PBMs, either fitted or expert, describe the response of ecophysiological processes to a wide range of environmental conditions, they can provide a better estimate of the environmental conditions in which species could have survived 12000 years ago, under climates much more dissimilar to present conditions. A potential limitation of our approach though is that we cannot account for very rare and really long-distance dispersion events, as well as the influence of humans. For example all models failed to predict deciduous *Quercus* in the British Isles before the Early Holocene sea level rise and the opening of the Strait of Dover (Fig. 2; 77), even though the land-sea mask changed throughout the simulations. It remains unclear whether this failure is due to the migration models’ misrepresentation of very long-distance dispersion events of seeds (e.g. by humans or jays, across major dispersal barrier), or a consistent misprediction by both CSDMs and PBMs of more northern refugia.

The recent efforts to gather fossil pollen data and make them openly available (61) allow us to objectively assess model performance under climate conditions vastly different from those used for their calibration. From 11.5 kyr BP onwards, climate dissimilarity varies between 0.29 and 0.08, a level equivalent to what we might experience in the second quarter of the 21st century (Fig. 1). The consistency of model projections with past observations does not demonstrate that model projections will be valid in the future (78), but making such comparisons allows to make a critical step towards enhancing our understanding of model transferability. As more and more pollen data becomes available, we could cover a wider range of conditions, notably prior 11.5 kyr BP. Our simulations nevertheless started at 12 kyr BP, when climatic dissimilarity was at its highest, and transitioned rapidly to a climate more analogous to historical state. The uncertainties on the initial conditions had thus a significant influence on the simulation outcomes. In the future, on the contrary, uncertainties on the initial conditions will be much lower as models will start from the known distributions of species, and uncertainties will increase as simulations proceed towards increasingly dissimilar climatic conditions, especially since these conditions will extent beyond the range experienced in the past (Figure S3).

While quantifying the uncertainty in model projections remains challenging, our results pave the way for drastic improvement in model evaluation. The discrepancies between model performances we observed highlights the importance of considering various modelling methods to capture the full range of uncertainties associated with future projections. It implies that we should not rely solely on the model’s own prediction dispersion to estimate projection uncertainties, nor on very similar modelling approaches, especially when climate dissimilarity sharply increases. Moreover, models will have to consider that tree colonization dynamics will likely be very different in the future because it will not only occur from a few refugia but from wider continuous ranges, and direct anthropogenic factors, such as sylvicultural practices and assisted species migration, will also shape the composition of forests (79).

Fitted PBMs bring together the strengths from both CSDMs and expert PBMs approaches by describing causal relationships between environmental conditions and species performance (i.e. from process-based approaches) and precise estimates of parameter values (from correlative approaches). The differences between expert and fitted PBMs in the Middle-Late Holocene pinpoint some issues in expert parameterization that requires to combine various methods to cope with both the scarcity of data for each ecophysiological process modelled and sometimes non-measurable parameters (e.g. 80). Some parameters in these relations can be measured directly, and exhibit little variability across a species range (e.g. water potential leading to 50% of vessels embolism). However, the measurement of parameters in controlled conditions does not necessarily guarantee their external validity *in natura* (81) where numerous factors, not represented in laboratory conditions, can also affect the process modelled (but see (82)). Other parameters are in addition either highly variable because of local adaptation over long period, difficult-to-measure or so far unmeasurable (e.g. bud dormancy). Therefore, expert PBMs can suffer from uncertainties entailed in the measurements of some of their parameters, and from spurious data specific to few locations which do not represent sufficiently well all the conditions the species can experience all over its range. For these reasons, inverse calibration presents a valuable opportunity to estimate the values of PBM parameters especially difficult to estimate otherwise (23, 83). However, inverse calibration does not warranty the correct estimation of parameter values and needs to be used critically and with caution (40).

Our unique multi-model comparison across the Holocene demonstrates that our understanding of biological mechanisms embedded into process-based models represent a real advantage over the empirical relationships used in CSDMs to increase projections reliability for the coming decades. However, data availability limits our ability to parameterize these models, and could explain the difficulty to use them more widely for global impact studies. Fitted PBMs may overcome this problem by using more data at a larger geographical scale, while keeping the predictive strength of causal relationships. Given ongoing improvements in computational methods and the availability of new global-scale measurements (e.g. forest structure and growth with remote sensing and LiDAR data), extensive calibration and more widespread application of process-based models seems now possible as well as an increase in model projections reliability.

## ACKNOWLEDGEMENTS

The authors are deeply grateful for many helpful comments from Elizabeth M. Wolkovich, which have greatly enriched this manuscript. They would also like to thank Christophe Randin for his valuable input throughout the course of this research, and Jed O. Kaplan for making GWGEN and LPJ-LMfire codes openly available. They acknowledge the support and computing resources of GenOuest and TGCC-CEA.

Future climate scenarios used were from the NEX-GDDP-CMIP6 dataset, prepared by the Climate Analytics Group and NASA Ames Research Center using the NASA Earth Exchange and distributed by the NASA Center for Climate Simulation (NCCS). They acknowledge the World Climate Research Programme, which, through its Working Group on Coupled Modeling, coordinated and promoted CMIP6. V.V. was supported by a PhD Fellowship from the GAIA doctoral school of the University of Montpellier, France.

## Bibliography

1. Terence P. Dawson, Stephen T. Jackson, Joanna I. House, Iain Colin Prentice, and Georgina M. Mace. Beyond Predictions: Biodiversity Conservation in a Changing Climate. Science, 332(6025):53–58, April 2011. doi: 10.1126/science.1200303.

2. Nicolas Mouquet, Yvan Lagadeuc, Vincent Devictor, Luc Doyen, Anne Duputié, Damien Eveillard, Denis Faure, Eric Garnier, Olivier Gimenez, Philippe Huneman, Franck Jabot, Philippe Jarne, Dominique Joly, Romain Julliard, Sonia Kéfi, Gael J. Kergoat, Sandra Lavorel, Line Le Gall, Laurence Meslin, Serge Morand, Xavier Morin, Hélène Morlon, Gilles Pinay, Roger Pradel, Frank M. Schurr, Wilfried Thuiller, and Michel Loreau. REVIEW: Predictive ecology in a changing world. Journal of Applied Ecology, 52(5):1293–1310, 2015. ISSN 1365-2664. doi: 10.1111/1365-2664.12482.

3. Michela Pacifici, Wendy B. Foden, Piero Visconti, James E. M. Watson, Stuart H. M. Butchart, Kit M. Kovacs, Brett R. Scheffers, David G. Hole, Tara G. Martin, H. Resit Akçakaya, Richard T. Corlett, Brian Huntley, David Bickford, Jamie A. Carr, Ary A. Hoffmann, Guy F. Midgley, Paul Pearce-Kelly, Richard G. Pearson, Stephen E. Williams, Stephen G. Willis, Bruce Young, and Carlo Rondinini. Assessing species vulnerability to climate change. Nature Climate Change, 5(3):215–224, March 2015. ISSN 1758-6798. doi: 10.1038/nclimate2448.

4. P. J. Bartlein, S. P. Harrison, S. Brewer, S. Connor, B. A. S. Davis, K. Gajewski, J. Guiot, T. I. Harrison-Prentice, A. Henderson, O. Peyron, I. C. Prentice, M. Scholze, H. Seppä, B. Shuman, S. Sugita, R. S. Thompson, A. E. Viau, J. Williams, and H. Wu. Pollen-based continental climate reconstructions at 6 and 21ka: a global synthesis. Climate Dynamics, 37(3):775–802, August 2011. ISSN 1432-0894. doi: 10.1007/s00382-010-0904-1.

5. Damien A. Fordham, Stephen T. Jackson, Stuart C. Brown, Brian Huntley, Barry W. Brook, Dorthe Dahl-Jensen, M. Thomas P. Gilbert, Bette L. Otto-Bliesner, Anders Svensson, Spyros Theodoridis, Janet M. Wilmshurst, Jessie C. Buettel, Elisabetta Canteri, Matthew McDowell, Ludovic Orlando, July A. Pilowsky, Carsten Rahbek, and David Nogues-Bravo. Using paleo-archives to safeguard biodiversity under climate change. Science, 369(6507): eabc5654, August 2020. doi: 10.1126/science.abc5654.

6. David E. Uribe-Rivera, Gurutzeta Guillera-Arroita, Saras M. Windecker, Patricio Pliscoff, and Brendan A. Wintle. The predictive performance of process-explicit range change models remains largely untested. Ecography, 2023(4):e06048, 2023. doi: 10.1111/ecog.06048.

7. Pascale Braconnot, Sandy P. Harrison, Masa Kageyama, Patrick J. Bartlein, Valerie Masson-Delmotte, Ayako Abe-Ouchi, Bette Otto-Bliesner, and Yan Zhao. Evaluation of climate models using palaeoclimatic data. Nature Climate Change, 2(6):417–424, June 2012. ISSN 1758-6798. doi: 10.1038/nclimate1456.

8. Kaitlin C. Maguire, Diego Nieto-Lugilde, Matthew C. Fitzpatrick, John W. Williams, and Jessica L. Blois. Modeling species and community responses to past, present, and future episodes of climatic and ecological change. Annual Review of Ecology, Evolution, and Systematics, 46(1):343–368, 2015. doi: 10.1146/annurev-ecolsys-112414-054441.

9. Samuel D. Veloz, John W. Williams, Jessica L. Blois, Feng He, Bette Otto-Bliesner, and Zhengyu Liu. No-analog climates and shifting realized niches during the late quaternary: implications for 21st-century predictions by species distribution models. Global Change Biology, 18(5):1698–1713, 2012. ISSN 1365-2486. doi: 10.1111/j.1365-2486.2011.02635.x.

10. Peter B. Pearman, Christophe F. Randin, Olivier Broennimann, Pascal Vittoz, Willem O. van der Knaap, Robin Engler, Gwenaelle Le Lay, Niklaus E. Zimmermann, and Antoine Guisan. Prediction of plant species distributions across six millennia. Ecology Letters, 11 (4):357–369, 2008. ISSN 1461-0248. doi: 10.1111/j.1461-0248.2007.01150.x.

11. David R. Roberts and Andreas Hamann. Predicting potential climate change impacts with bioclimate envelope models: a palaeoecological perspective. Global Ecology and Biogeography, 21(2):121–133, 2012. ISSN 1466-8238. doi: 10.1111/j.1466-8238.2011.00657.x.

12. A. M. Foley, D. Dalmonech, A. D. Friend, F. Aires, A. T. Archibald, P. Bartlein, L. Bopp, J. Chappellaz, P. Cox, N. R. Edwards, G. Feulner, P. Friedlingstein, S. P. Harrison, P. O. Hopcroft, C. D. Jones, J. Kolassa, J. G. Levine, I. C. Prentice, J. Pyle, N. Vázquez Riveiros, E. W. Wolff, and S. Zaehle. Evaluation of biospheric components in earth system models using modern and palaeo-observations: the state-of-the-art. Biogeosciences, 10(12):8305–8328, 2013. doi: 10.5194/bg-10-8305-2013.

13. Kaitlin C. Maguire, Diego Nieto-Lugilde, Jessica L. Blois, Matthew C. Fitzpatrick, John W. Williams, Simon Ferrier, and David J. Lorenz. Controlled comparison of species- and community-level models across novel climates and communities. Proceedings of the Royal Society B: Biological Sciences, 283(1826):20152817, March 2016. doi: 10.1098/rspb.2015.2817.

14. John W. Williams, Stephen T. Jackson, and John E. Kutzbach. Projected distributions of novel and disappearing climates by 2100 AD. Proceedings of the National Academy of Sciences, 104(14):5738–5742, April 2007. ISSN 0027-8424, 1091-6490. doi: 10.1073/pnas.0606292104.

15. Matthew C. Fitzpatrick, Jessica L. Blois, John W. Williams, Diego Nieto-Lugilde, Kaitlin C. Maguire, and David J. Lorenz. How will climate novelty influence ecological forecasts? Using the Quaternary to assess future reliability. Global Change Biology, 24(8):3575–3586, 2018. ISSN 1365-2486. doi: 10.1111/gcb.14138.

16. K. D. Burke, J. W. Williams, M. A. Chandler, A. M. Haywood, D. J. Lunt, and B. L. Otto-Bliesner. Pliocene and Eocene provide best analogs for near-future climates. Proceedings of the National Academy of Sciences, 115(52):13288–13293, December 2018. ISSN 0027-8424, 1091-6490. doi: 10.1073/pnas.1809600115.

17. Peter U. Clark, Arthur S. Dyke, Jeremy D. Shakun, Anders E. Carlson, Jorie Clark, Barbara Wohlfarth, Jerry X. Mitrovica, Steven W. Hostetler, and A. Marshall McCabe. The last glacial maximum. Science, 325(5941):710–714, 2009. doi: 10.1126/science.1172873.

18. Carsten F. Dormann, Stanislaus J. Schymanski, Juliano Cabral, Isabelle Chuine, Catherine Graham, Florian Hartig, Michael Kearney, Xavier Morin, Christine Römermann, Boris Schröder, and Alexander Singer. Correlation and process in species distribution models: bridging a dichotomy. Journal of Biogeography, 39(12):2119–2131, 2012. ISSN 1365-2699. doi: 10.1111/j.1365-2699.2011.02659.x.

19. Marc Hanewinkel, Dominik A. Cullmann, Mart-Jan Schelhaas, Gert-Jan Nabuurs, and Niklaus E. Zimmermann. Climate change may cause severe loss in the economic value of European forest land. Nature Climate Change, 3(3):203–207, March 2013. ISSN 1758-6798. doi: 10.1038/nclimate1687.

20. M. Lukas Seehausen, Jacques Régnière, Véronique Martel, and Sandy M. Smith. Developmental and reproductive responses of the spruce budworm (Lepidoptera: Tortricidae) parasitoid Tranosema rostrale (Hymenoptera: Ichneumonidae) to temperature. Journal of Insect Physiology, 98:38–46, April 2017. ISSN 0022-1910. doi: 10.1016/j.jinsphys.2016.11.008.

21. Mingkai Jiang, Belinda E. Medlyn, John E. Drake, Remko A. Duursma, Ian C. Anderson, Craig V. M. Barton, Matthias M. Boer, Yolima Carrillo, Laura Castañeda-Gómez, Luke Collins, Kristine Y. Crous, Martin G. De Kauwe, Bruna M. dos Santos, Kathryn M. Emmerson, Sarah L. Facey, Andrew N. Gherlenda, Teresa E. Gimeno, Shun Hasegawa, Scott N. Johnson, Astrid Kännaste, Catriona A. Macdonald, Kashif Mahmud, Ben D. Moore, Loïc Nazaries, Elizabeth H. J. Neilson, Uffe N. Nielsen, Ülo Niinemets, Nam Jin Noh, Raúl Ochoa-Hueso, Varsha S. Pathare, Elise Pendall, Johanna Pihlblad, Juan Piñeiro, Jeff R. Powell, Sally A. Power, Peter B. Reich, Alexandre A. Renchon, Markus Riegler, Riikka Rinnan, Paul D. Rymer, Roberto L. Salomón, Brajesh K. Singh, Benjamin Smith, Mark G. Tjoelker, Jennifer K. M. Walker, Agnieszka Wujeska-Klause, Jinyan Yang, Sönke Zaehle, and David S. Ellsworth. The fate of carbon in a mature forest under carbon dioxide enrichment. Nature, 580(7802):227–231, April 2020. ISSN 1476-4687. doi: 10.1038/s41586-020-2128-9.

22. Jordane Gavinet, Jean-Marc Ourcival, and Jean-Marc Limousin. Rainfall exclusion and thinning can alter the relationships between forest functioning and drought. New Phytologist, 223(3):1267–1279, 2019. ISSN 1469-8137. doi: 10.1111/nph.15860.

23. Margaret E. K. Evans, Cory Merow, Sydne Record, Sean M. McMahon, and Brian J. Enquist. Towards Process-based Range Modeling of Many Species. Trends in Ecology & Evolution, 31(11):860–871, November 2016. ISSN 0169-5347. doi: 10.1016/j.tree.2016.08.005.

24. Matthew R. Evans. Modelling ecological systems in a changing world. Philosophical Transactions of the Royal Society B: Biological Sciences, 367(1586):181–190, January 2012. doi: 10.1098/rstb.2011.0172.

25. Alexander Singer, Karin Johst, Thomas Banitz, Mike S. Fowler, Jürgen Groeneveld, Alvaro G. Gutiérrez, Florian Hartig, Rainer M. Krug, Matthias Liess, Glenn Matlack, Katrin M. Meyer, Guy Pe’er, Viktoriia Radchuk, Ana-Johanna Voinopol-Sassu, and Justin M. J. Travis. Community dynamics under environmental change: How can next generation mechanistic models improve projections of species distributions? Ecological Modelling, 326:63–74, April 2016. ISSN 0304-3800. doi: 10.1016/j.ecolmodel.2015.11.007.

26. Sean R. Connolly, Sally A. Keith, Robert K. Colwell, and Carsten Rahbek. Process, Mechanism, and Modeling in Macroecology. Trends in Ecology & Evolution, 32(11):835–844, November 2017. ISSN 0169-5347. doi: 10.1016/j.tree.2017.08.011.

27. M. C. Urban, G. Bocedi, A. P. Hendry, J.-B. Mihoub, G. Pe’er, A. Singer, J. R. Bridle, L. G. Crozier, L. De Meester, W. Godsoe, A. Gonzalez, J. J. Hellmann, R. D. Holt, A. Huth, K. Johst, C. B. Krug, P. W. Leadley, S. C. F. Palmer, J. H. Pantel, A. Schmitz, P. A. Zollner, and J. M. J. Travis. Improving the forecast for biodiversity under climate change. Science, 353(6304):aad8466, September 2016. doi: 10.1126/science.aad8466.

28. July A. Pilowsky, Robert K. Colwell, Carsten Rahbek, and Damien A. Fordham. Process-explicit models reveal the structure and dynamics of biodiversity patterns. Science Advances, 8(31):eabj2271, August 2022. doi: 10.1126/sciadv.abj2271.

29. Xavier Morin and Wilfried Thuiller. Comparing niche- and process-based models to reduce prediction uncertainty in species range shifts under climate change. Ecology, 90(5):1301–1313, 2009. ISSN 1939-9170. doi: 10.1890/08-0134.1.

30. Alissar Cheaib, Vincent Badeau, Julien Boe, Isabelle Chuine, Christine Delire, Eric Dufrêne, Christophe François, Emmanuel S. Gritti, Myriam Legay, Christian Pagé, Wilfried Thuiller, Nicolas Viovy, and Paul Leadley. Climate change impacts on tree ranges: model intercomparison facilitates understanding and quantification of uncertainty. Ecology Letters, 15(6): 533–544, 2012. ISSN 1461-0248. doi: 10.1111/j.1461-0248.2012.01764.x.

31. Emmanuel S. Gritti, Anne Duputié, Francois Massol, and Isabelle Chuine. Estimating consensus and associated uncertainty between inherently different species distribution models. Methods in Ecology and Evolution, 4(5):442–452, 2013. ISSN 2041-210X. doi: 10.1111/2041-210X.12032.

32. Damaris Zurell, Wilfried Thuiller, Jörn Pagel, Juliano S. Cabral, Tamara Münkemüller, Dominique Gravel, Stefan Dullinger, Signe Normand, Katja H. Schiffers, Kara A. Moore, and Niklaus E. Zimmermann. Benchmarking novel approaches for modelling species range dynamics. Global Change Biology, 22(8):2651–2664, 2016. ISSN 1365-2486. doi: 10.1111/gcb.13251.

33. Steven I. Higgins, Matthew J. Larcombe, Nicholas J. Beeton, Timo Conradi, and Henning Nottebrock. Predictive ability of a process-based versus a correlative species distribution model. Ecology and Evolution, 10(20):11043–11054, 2020. ISSN 2045-7758. doi: 10.1002/ece3.6712.

34. Damien A. Fordham, Cleo Bertelsmeier, Barry W. Brook, Regan Early, Dora Neto, Stuart C. Brown, Sébastien Ollier, and Miguel B. Araújo. How complex should models be? Comparing correlative and mechanistic range dynamics models. Global Change Biology, 24(3):1357–1370, 2018. ISSN 1365-2486. doi: 10.1111/gcb.13935.

35. Frédérik Saltré, Rémi Saint-Amant, Emmanuel S. Gritti, Simon Brewer, Cédric Gaucherel, Basil A. S. Davis, and Isabelle Chuine. Climate or migration: what limited European beech post-glacial colonization? Global Ecology and Biogeography, 22(11):1217–1227, 2013. ISSN 1466-8238. doi: 10.1111/geb.12085.

36. Melanie Ruosch, Renato Spahni, Fortunat Joos, Paul D. Henne, Willem O. van der Knaap, and Willy Tinner. Past and future evolution of Abies alba forests in Europe – comparison of a dynamic vegetation model with palaeo data and observations. Global Change Biology, 22 (2):727–740, 2016. ISSN 1365-2486. doi: 10.1111/gcb.13075.

37. Christoph Schwörer, Paul D. Henne, and Willy Tinner. A model-data comparison of Holocene timberline changes in the Swiss Alps reveals past and future drivers of mountain forest dynamics. Global Change Biology, 20(5):1512–1526, 2014. ISSN 1365-2486. doi: 10.1111/gcb.12456.

38. Natalie J. Briscoe, Jane Elith, Roberto Salguero-Gómez, José J. Lahoz-Monfort, James S. Camac, Katherine M. Giljohann, Matthew H. Holden, Bronwyn A. Hradsky, Michael R. Kearney, Sean M. McMahon, Ben L. Phillips, Tracey J. Regan, Jonathan R. Rhodes, Peter A. Vesk, Brendan A. Wintle, Jian D.L. Yen, and Gurutzeta Guillera-Arroita. Forecasting species range dynamics with process-explicit models: matching methods to applications. Ecology Letters, 22(11):1940–1956, 2019. ISSN 1461-0248. doi: 10.1111/ele.13348.

39. Edward Armstrong, Peter O. Hopcroft, and Paul J. Valdes. A simulated Northern Hemisphere terrestrial climate dataset for the past 60,000 years. Scientific Data, 6(1):265, November 2019. ISSN 2052-4463. doi: 10.1038/s41597-019-0277-1.

40. Victor Van der Meersch and Isabelle Chuine. Estimating process-based model parameters from species distribution data using the evolutionary algorithm CMA-ES. Methods in Ecology and Evolution, 14(7):1808–1820, 2023. ISSN 2041-210X. doi: 10.1111/2041-210X.14119.

41. Isabelle Chuine and Elisabeth G. Beaubien. Phenology is a major determinant of tree species range. Ecology Letters, 4(5):500–510, 2001. ISSN 1461-0248. doi: 10.1046/j.1461-0248.2001.00261.x.

42. Xavier Morin, Carol Augspurger, and Isabelle Chuine. Process-Based Modeling of Species’ Distributions: What Limits Temperate Tree Species’ Range Boundaries? Ecology, 88(9): 2280–2291, 2007. ISSN 1939-9170. doi: 10.1890/06-1591.1.

43. Anne Duputié, Alexis Rutschmann, Ophélie Ronce, and Isabelle Chuine. Phenological plasticity will not help all species adapt to climate change. Global Change Biology, 21(8):3062–3073, 2015. ISSN 1365-2486. doi: 10.1111/gcb.12914.

44. Julie Gauzere, Bertrand Teuf, Hendrik Davi, Luis-Miguel Chevin, Thomas Caignard, Bérangère Leys, Sylvain Delzon, Ophélie Ronce, and Isabelle Chuine. Where is the optimum? Predicting the variation of selection along climatic gradients and the adaptive value of plasticity. A case study on tree phenology. Evolution Letters, 4(2):109–123, 2020. ISSN 2056-3744. doi: 10.1002/evl3.160.

45. E. Dufrêne, H. Davi, C. François, G. le Maire, V. Le Dantec, and A. Granier. Modelling carbon and water cycles in a beech forest: Part I: Model description and uncertainty analysis on modelled NEE. Ecological Modelling, 185(2):407–436, July 2005. ISSN 0304-3800. doi: 10.1016/j.ecolmodel.2005.01.004.

46. H. Davi, E. Dufrêne, C. Francois, G. Le Maire, D. Loustau, A. Bosc, S. Rambal, A. Granier, and E. Moors. Sensitivity of water and carbon fluxes to climate changes from 1960 to 2100 in European forest ecosystems. Agricultural and Forest Meteorology, 141(1):35–56, December 2006. ISSN 0168-1923. doi: 10.1016/j.agrformet.2006.09.003.

47. N. Delpierre, K. Soudani, C. François, G. Le Maire, C. Bernhofer, W. Kutsch, L. Misson, S. Rambal, T. Vesala, and E. Dufrêne. Quantifying the influence of climate and biological drivers on the interannual variability of carbon exchanges in European forests through process-based modelling. Agricultural and Forest Meteorology, 154-155:99–112, March 2012. ISSN 0168-1923. doi: 10.1016/j.agrformet.2011.10.010.

48. Hendrik Davi and Maxime Cailleret. Assessing drought-driven mortality trees with physiological process-based models. Agricultural and Forest Meteorology, 232:279–290, January 2017. ISSN 0168-1923. doi: 10.1016/j.agrformet.2016.08.019.

49. Nikolaus Hansen and Andreas Ostermeier. Completely Derandomized Self-Adaptation in Evolution Strategies. Evolutionary Computation, 9(2):159–195, June 2001. ISSN 1063-6560. doi: 10.1162/106365601750190398.

50. Roozbeh Valavi, Gurutzeta Guillera-Arroita, José J. Lahoz-Monfort, and Jane Elith. Predictive performance of presence-only species distribution models: a benchmark study with reproducible code. Ecological Monographs, 92(1):e01486, 2022. ISSN 1557-7015. doi: 10.1002/ecm.1486.

51. Joaquín Muñoz-Sabater, Emanuel Dutra, Anna Agustí-Panareda, Clément Albergel, Gabriele Arduini, Gianpaolo Balsamo, Souhail Boussetta, Margarita Choulga, Shaun Harrigan, and Hans Hersbach. ERA5-Land: A state-of-the-art global reanalysis dataset for land applications. Earth system science data, 13(9):4349–4383, 2021.

52. Brigitta Tóth, Melanie Weynants, László Pásztor, and Tomislav Hengl. 3D soil hydraulic database of Europe at 250 m resolution. Hydrological Processes, 31(14):2662–2666, 2017. ISSN 1099-1085. doi: 10.1002/hyp.11203.

53. Tomislav Hengl, Jorge Mendes de Jesus, Gerard B. M. Heuvelink, Maria Ruiperez Gonzalez, Milan Kilibarda, Aleksandar Blagotic, Wei Shangguan, Marvin N. Wright, Xiaoyuan Geng, Bernhard Bauer-Marschallinger, Mario Antonio Guevara, Rodrigo Vargas, Robert A. MacMillan, Niels H. Batjes, Johan G. B. Leenaars, Eloi Ribeiro, Ichsani Wheeler, Stephan Mantel, and Bas Kempen. SoilGrids250m: Global gridded soil information based on machine learning. PLOS ONE, 12(2):e0169748, February 2017. ISSN 1932-6203. doi: 10.1371/journal.pone.0169748.

54. Achille Mauri, Giovanni Strona, and Jesús San-Miguel-Ayanz. EU-Forest, a high-resolution tree occurrence dataset for Europe. Scientific Data, 4(1):160123, January 2017. ISSN 2052-4463. doi: 10.1038/sdata.2016.123.

55. David R. Roberts, Volker Bahn, Simone Ciuti, Mark S. Boyce, Jane Elith, Gurutzeta Guillera-Arroita, Severin Hauenstein, José J. Lahoz-Monfort, Boris Schröder, Wilfried Thuiller, David I. Warton, Brendan A. Wintle, Florian Hartig, and Carsten F. Dormann. Cross-validation strategies for data with temporal, spatial, hierarchical, or phylogenetic structure. Ecography, 40(8):913–929, 2017. ISSN 1600-0587. doi: 10.1111/ecog.02881.

56. W. R. Peltier, D. F. Argus, and R. Drummond. Space geodesy constrains ice age terminal deglaciation: The global ICE-6G_c (VM5a) model. Journal of Geophysical Research: Solid Earth, 120(1):450–487, 2015. ISSN 2169-9356. doi: 10.1002/2014JB011176.

57. P. S. Sommer and J. O. Kaplan. A globally calibrated scheme for generating daily meteorology from monthly statistics: Global-wgen (gwgen) v1.0. Geoscientific Model Development, 10(10):3771–3791, 2017. doi: 10.5194/gmd-10-3771-2017.

58. M. Pfeiffer, A. Spessa, and J. O. Kaplan. A model for global biomass burning in preindustrial time: Lpj-lmfire (v1.0). Geoscientific Model Development, 6(3):643–685, 2013. doi: 10.5194/gmd-6-643-2013.

59. Richard Allen, Luis Pereira, Dirk Raes, and Martin Smith. Crop evapotranspiration: Guidelines for computing crop water requirements. FAO Irrigation and Drainage Papers, 56, 1998.

60. U. Herzschuh, C. Li, T. Böhmer, A. K. Postl, B. Heim, A. A. Andreev, X. Cao, M. Wieczorek, and J. Ni. Legacypollen 1.0: a taxonomically harmonized global late quaternary pollen dataset of 2831 records with standardized chronologies. Earth System Science Data, 14 (7):3213–3227, 2022. doi: 10.5194/essd-14-3213-2022.

61. John W. Williams, Eric C. Grimm, Jessica L. Blois, Donald F. Charles, Edward B. Davis, Simon J. Goring, Russell W. Graham, Alison J. Smith, Michael Anderson, Joaquin Arroyo-Cabrales, Allan C. Ashworth, Julio L. Betancourt, Brian W. Bills, Robert K. Booth, Philip I. Buckland, B. Brandon Curry, Thomas Giesecke, Stephen T. Jackson, Claudio Latorre, Jonathan Nichols, Timshel Purdum, Robert E. Roth, Michael Stryker, and Hikaru Takahara. The Neotoma Paleoecology Database, a multiproxy, international, community-curated data resource. Quaternary Research, 89(1):156–177, January 2018. ISSN 0033-5894, 1096-0287. doi: 10.1017/qua.2017.105.

62. John W. Williams, Robert L. Summers, and Thompson Webb III. Applying plant functional types to construct biome maps from eastern North American pollen data: comparisons with model results. Quaternary Science Reviews, 17(6):607–627, April 1998. ISSN 0277-3791. doi: 10.1016/S0277-3791(98)00014-6.

63. J. Jalas and J. Suominen. Atlas Florae Europaeae. Committee for Mapping the Flora of Europe and Societas Biologica Fennica Vanamo, Helsinki, Finland, 1972–2005.

64. Udo Bohn, Robert Neuhäusl, Gisela Gollub, Christoph Hettwer, Zdenka Neuhäuslová, Thomas Raus, Heinz Schlüter, and Herbert Weber. Map of the natural vegetation of europe - scale 1:2500000, 2003.

65. Jens-Christian Svenning and Flemming Skov. Limited filling of the potential range in european tree species. Ecology Letters, 7(7):565–573, 2004. doi: 10.1111/j.1461-0248.2004.00614.x.

66. Robin Engler, Wim Hordijk, and Antoine Guisan. The MIGCLIM R package – seamless integration of dispersal constraints into projections of species distribution models. Ecography, 35(10):872–878, 2012. ISSN 1600-0587. doi: 10.1111/j.1600-0587.2012.07608.x.

67. Deborah Zani, Veiko Lehsten, and Heike Lischke. Tree migration in the dynamic, global vegetation model LPJ-GM 1.1: efficient uncertainty assessment and improved dispersal kernels of European trees. Geoscientific Model Development, 15(12):4913–4940, June 2022. ISSN 1991-959X. doi: 10.5194/gmd-15-4913-2022.

68. Boris Leroy, Robin Delsol, Bernard Hugueny, Christine N. Meynard, Chéïma Barhoumi, Morgane Barbet-Massin, and Céline Bellard. Without quality presence–absence data, discrimination metrics such as TSS can be misleading measures of model performance. Journal of Biogeography, 45(9):1994–2002, 2018. ISSN 1365-2699. doi: 10.1111/jbi.13402.

69. Ian Harris, Timothy J. Osborn, Phil Jones, and David Lister. Version 4 of the CRU TS monthly high-resolution gridded multivariate climate dataset. Scientific Data, 7(1):109, April 2020. ISSN 2052-4463. doi: 10.1038/s41597-020-0453-3.

70. Benjamin Blonder, Cecina Babich Morrow, Brian Maitner, David J. Harris, Christine Lamanna, Cyrille Violle, Brian J. Enquist, and Andrew J. Kerkhoff. New approaches for delineating n-dimensional hypervolumes. Methods in Ecology and Evolution, 9(2):305–319, 2018. ISSN 2041-210X. doi: 10.1111/2041-210X.12865.

71. Kevin D. Burke, John W. Williams, Simon Brewer, Walter Finsinger, Thomas Giesecke, David J. Lorenz, and Alejandro Ordonez. Differing climatic mechanisms control transient and accumulated vegetation novelty in Europe and eastern North America. Philosophical Transactions of the Royal Society B: Biological Sciences, 374(1788):20190218, December 2019. doi: 10.1098/rstb.2019.0218.

72. Bridget Thrasher, Weile Wang, Andrew Michaelis, Forrest Melton, Tsengdar Lee, and Ramakrishna Nemani. NASA Global Daily Downscaled Projections, CMIP6. Scientific Data, 9 (1):262, June 2022. ISSN 2052-4463. doi: 10.1038/s41597-022-01393-4.

73. Robert Kubinec. Ordered beta regression: A parsimonious, well-fitting model for continuous data with lower and upper bounds. Political Analysis, 31(4):519–536, 2023. doi: 10.1017/pan.2022.20.

74. Stan Development Team.RStan: the R interface to Stan, 2023. R package version 2.32.3.

75. Colin M. Beale and Jack J. Lennon. Incorporating uncertainty in predictive species distribution modelling. Philosophical Transactions of the Royal Society B: Biological Sciences, 367 (1586):247–258, January 2012. doi: 10.1098/rstb.2011.0178.

76. Frans Berkhout. Reconstructing boundaries and reason in the climate debate. Global Environmental Change, 20(4):565–569, October 2010. ISSN 0959-3780. doi: 10.1016/j.gloenvcha.2010.07.006.

77. D. E. Smith, S. Harrison, C. R. Firth, and J. T. Jordan. The early Holocene sea level rise. Quaternary Science Reviews, 30(15):1846–1860, July 2011. ISSN 0277-3791. doi: 10.1016/j.quascirev.2011.04.019.

78. Naomi Oreskes, Kristin Shrader-Frechette, and Kenneth Belitz. Verification, Validation, and Confirmation of Numerical Models in the Earth Sciences. Science, 263(5147):641–646, February 1994. doi: 10.1126/science.263.5147.641.

79. Sally N. Aitken and Jordan B. Bemmels. Time to get moving: assisted gene flow of forest trees. Evolutionary Applications, 9(1):271–290, 2016. ISSN 1752-4571. doi: 10.1111/eva.12293.

80. M. D. Cáceres, R. Molowny-Horas, A. Cabon, J. Martínez-Vilalta, M. Mencuccini, R. García-Valdés, D. Nadal-Sala, S. Sabaté, N. Martin-StPaul, X. Morin, F. D’Adamo, E. Batllori, and A. Améztegui. Medfate 2.9.3: a trait-enabled model to simulate mediterranean forest function and dynamics at regional scales. Geoscientific Model Development, 16(11):3165–3201, 2023. doi: 10.5194/gmd-16-3165-2023.

81. Daphné Asse Christophe F. Randin, Marc Bonhomme, Anne Delestrade, and Isabelle Chuine. Process-based models outcompete correlative models in projecting spring phenology of trees in a future warmer climate. Agricultural and Forest Meteorology, 285-286: 107931, May 2020. ISSN 0168-1923. doi: 10.1016/j.agrformet.2020.107931.

82. Akiko Satake, Tetsuhiro Kawagoe, Yukari Saburi, Yukako Chiba, Gen Sakurai, and Hiroshi Kudoh. Forecasting flowering phenology under climate warming by modelling the regulatory dynamics of flowering-time genes. Nature Communications, 4(1):2303, August 2013. ISSN 2041-1723. doi: 10.1038/ncomms3303.

83. F. Hartig, C. Dislich, T. Wiegand, and A. Huth. Technical Note: Approximate Bayesian parameterization of a process-based tropical forest model. Biogeosciences, 11(4):1261–1272, February 2014. ISSN 1726-4170. doi: 10.5194/bg-11-1261-2014.

