## Supplementary information for "Biological mechanisms are necessary to improve projections of species range shifts"

Victor Van der Meersch[1*], Edward Armstrong[2], Florent Mouillot[1], Anne Duputié[3], Hendrik Davi[4], Frédérik Saltré[5], Isabelle Chuine[1]

[1]CEFE, Université de Montpellier, CNRS, EPHE, IRD, Montpellier, France.
[2]Dept. of Geosciences and Geography, University of Helsinki, Helsinki, Finland.
[3]UMR 8198-EEP-Evo-Eco-Paleo, Université de Lille, CNRS, Lille, France.
[4]URFM, INRAE, Avignon, France.
[5]Global Ecology, College of Science and Engineering, Flinders University, Adelaide, Australia.

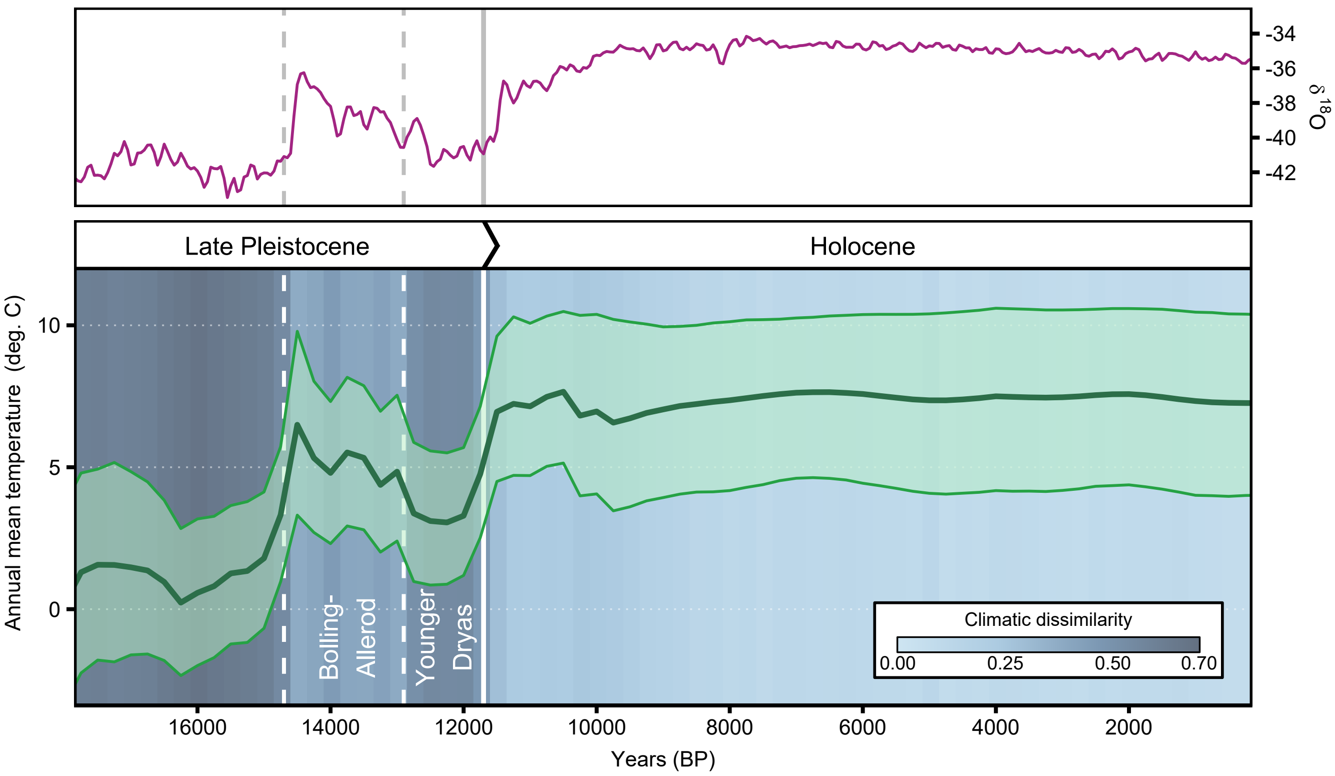


**Fig. 1 Chronology of past climate across the Late Pleistocene and the Holocene.** Upper panel shows the evolution of North Greenland Ice-Core Project (NGRIP) oxygen isotope 18 values (permille) as 50 year mean values [1]. Lower panel shows the average annual temperature across Europe, from HadCM3B simulations [2]. Shaded area represents interquartile range. Blue background represents the level of climatic dissimilarity (see Methods).

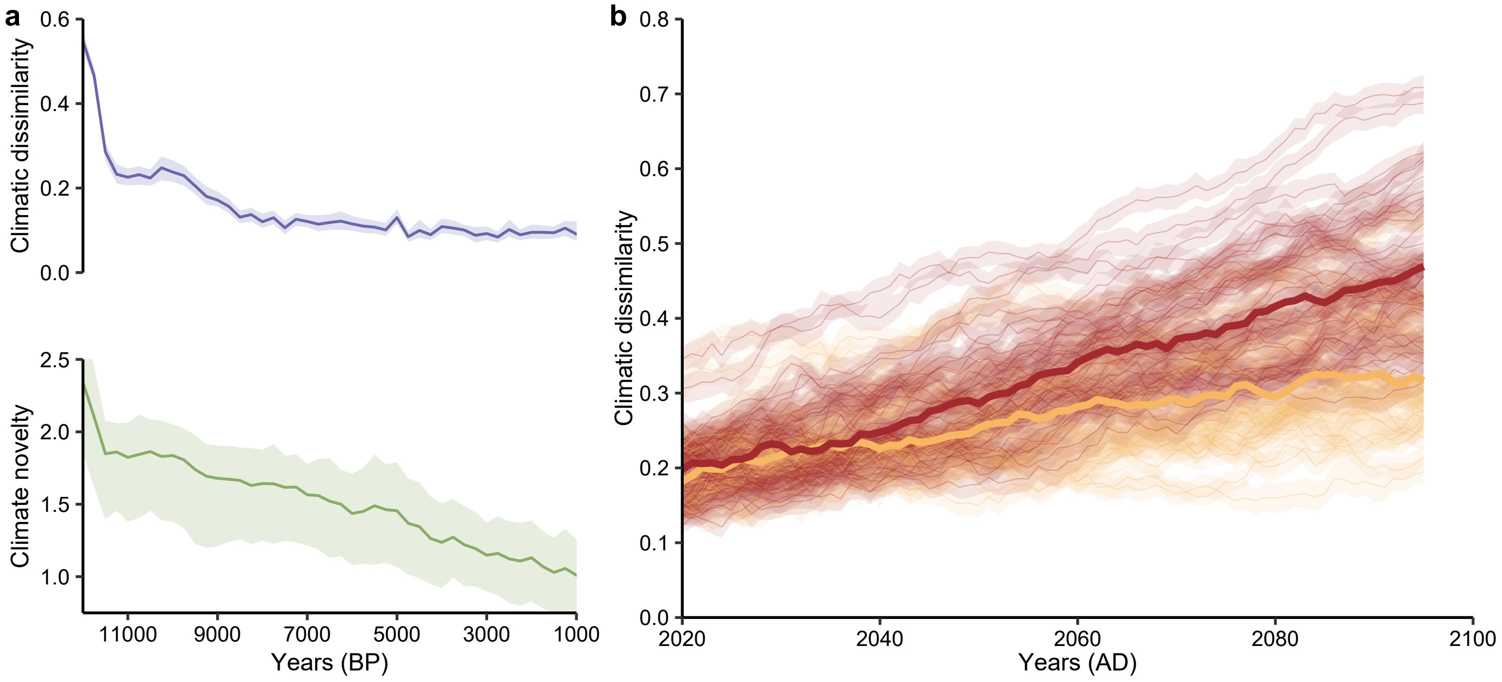


**Fig. 2** **(a)** Climatic dissimilarity as calculated in this paper and climate novelty calculated following [3]. The reference period is 1901-2000. **(b)** Evolution of future climatic dissimilarity, across the 34 GCM simulations [4]. Yellow and red correspond to SSP245 and SSP585 scenarios. Shaded areas represent 95% confidence interval for climatic dissimilarity/interquartile range for climate novelty.

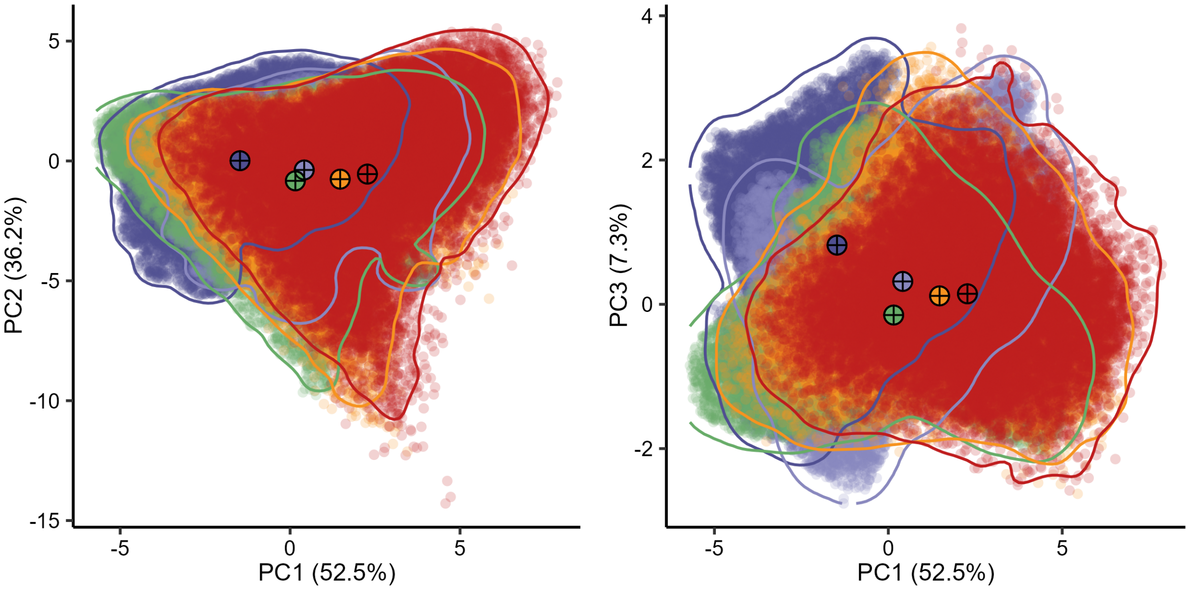


**Fig. 3** Climatic hypervolumes for Early Holocene period (dark and light blue, respectively 12000 BP and 11500 BP) and 2090 projections (orange and red, respectively SSP2 and SSP5 scenarios), as compared to historical period (green, 1901-2000). Hypervolume contours are generated using a two-dimensional kernel density estimation. Bigger points represent hypervolume centroids. Note that SSP2 and SSP5 points where randomly sampled in the 34 GCM hypervolumes used in this study [4]. PC1-3 correspond to the first three principal component axis from three-month means temperature and three-month sums of precipitation (see Methods), with the percentage of variation explained in parenthesis. Temperature contributes mostly to PC1, whereas precipitation contributes mostly to PC2 and PC3.

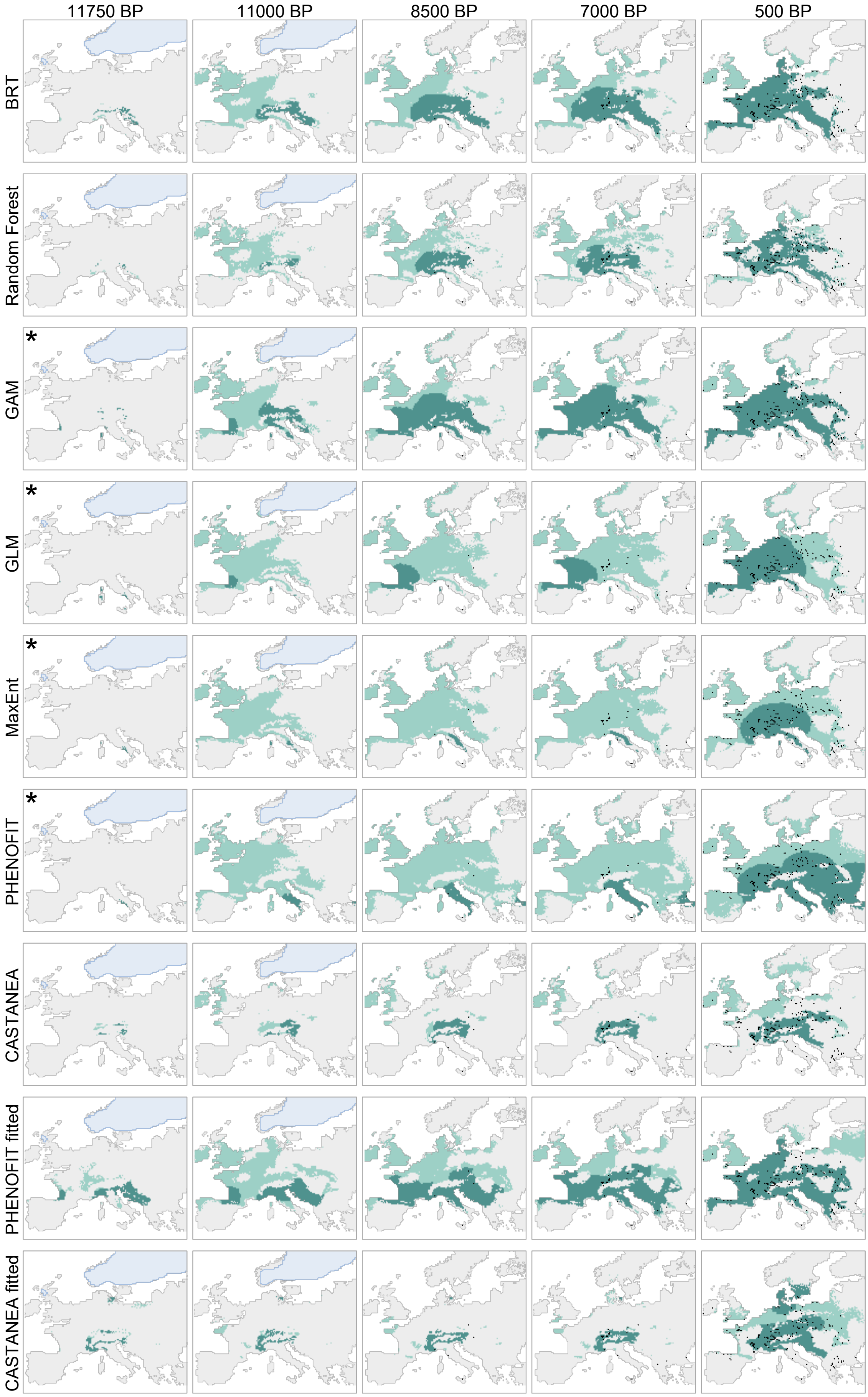

**Fig. 4** Same caption as Fig. 2 in the main text, for beech.

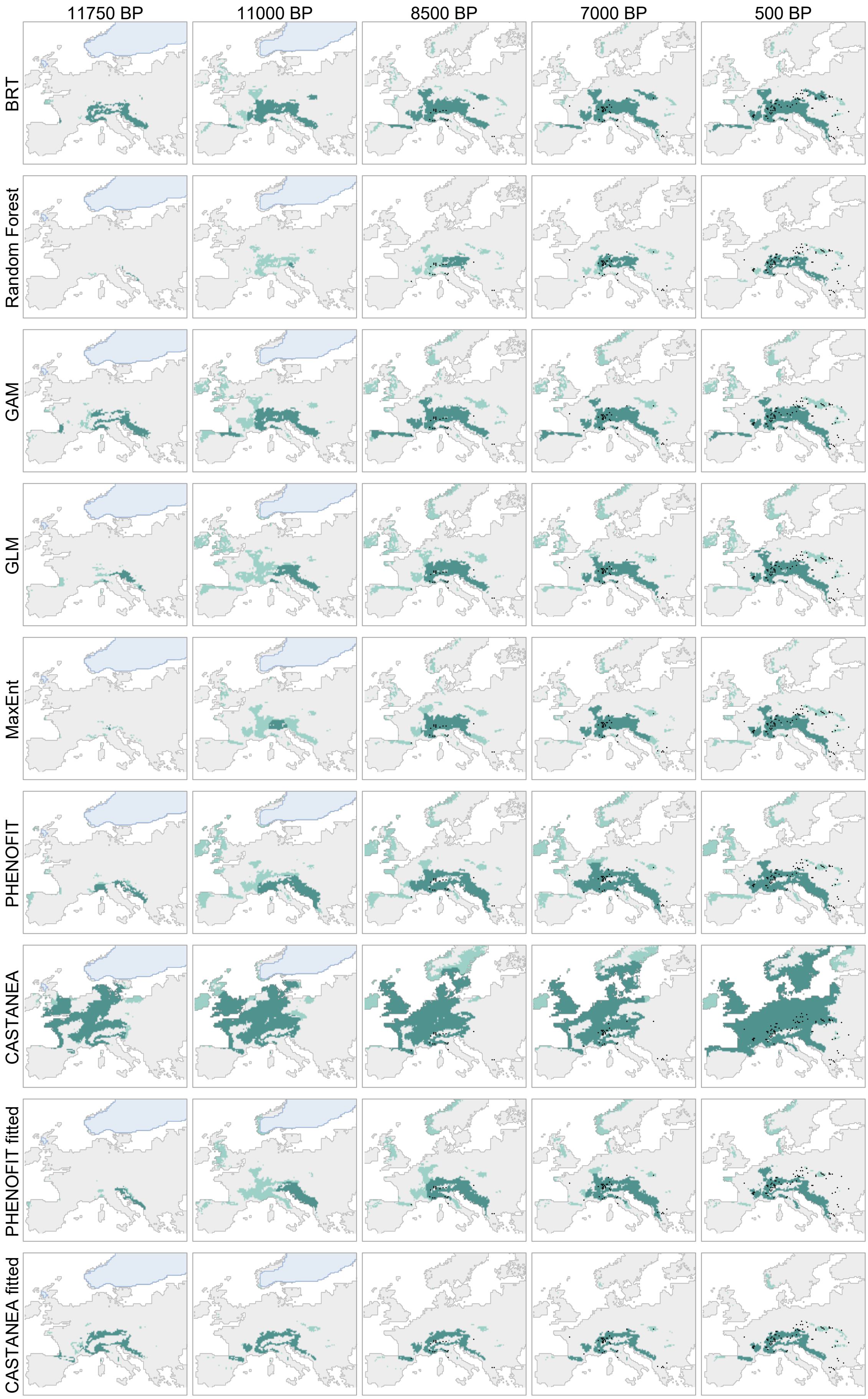


**Fig. 5** Same caption as Fig. 2 in the main text, for fir.

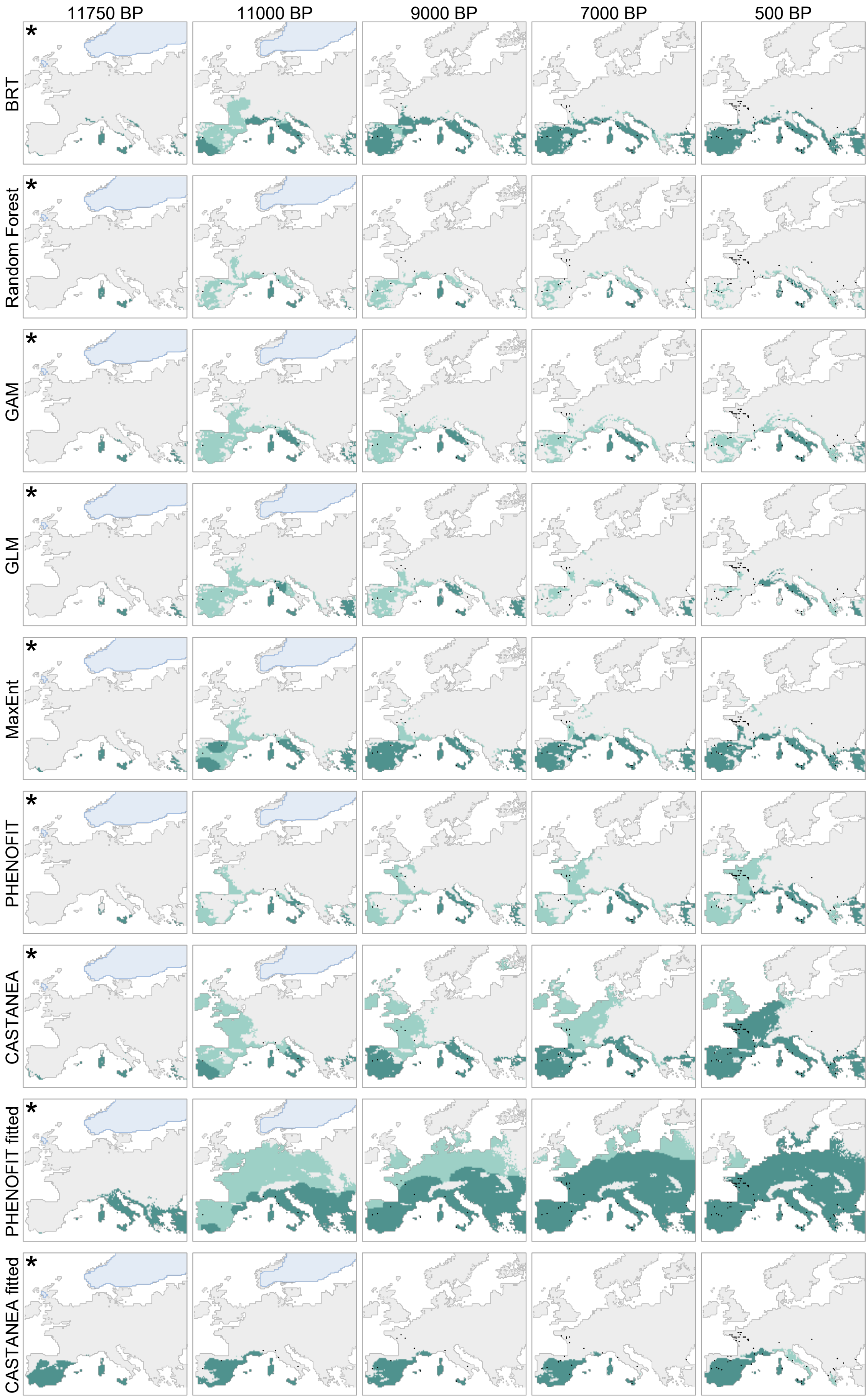


**Fig. 6** Same caption as Fig. 2 in the main text, for evergreen oak.

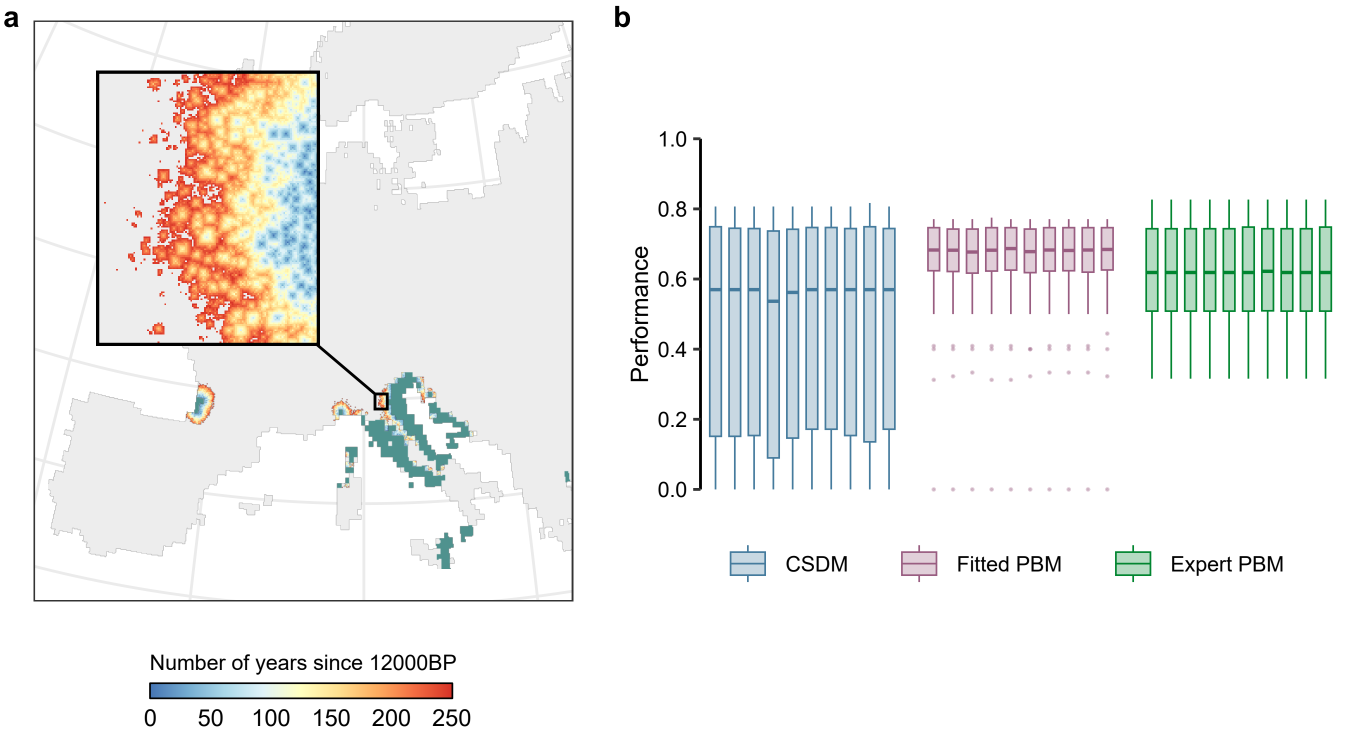


**Fig. 7** **(a)** Illustration of the first 250 years of deciduous oak migration process, with PHENOFIT fitted model. Dark green area represents migration starting points at 12000 BP. **(b)** Variation of model performance for deciduous *Quercus*, due to the stochasticity of migration simulations.

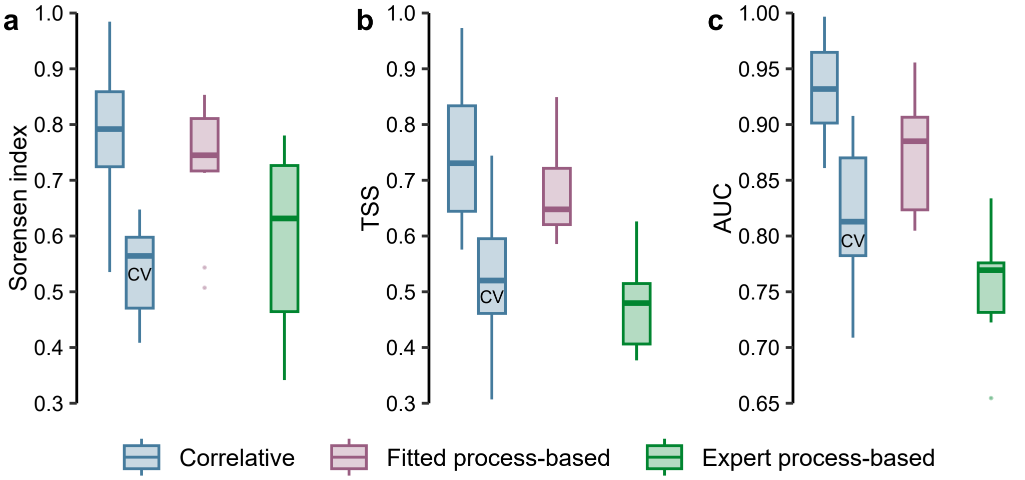


**Fig. 8** **Model performance in historical conditions.** "CV" stands for "cross-validation", when correlative extrapolation errors were assessed using a block cross-validation method.

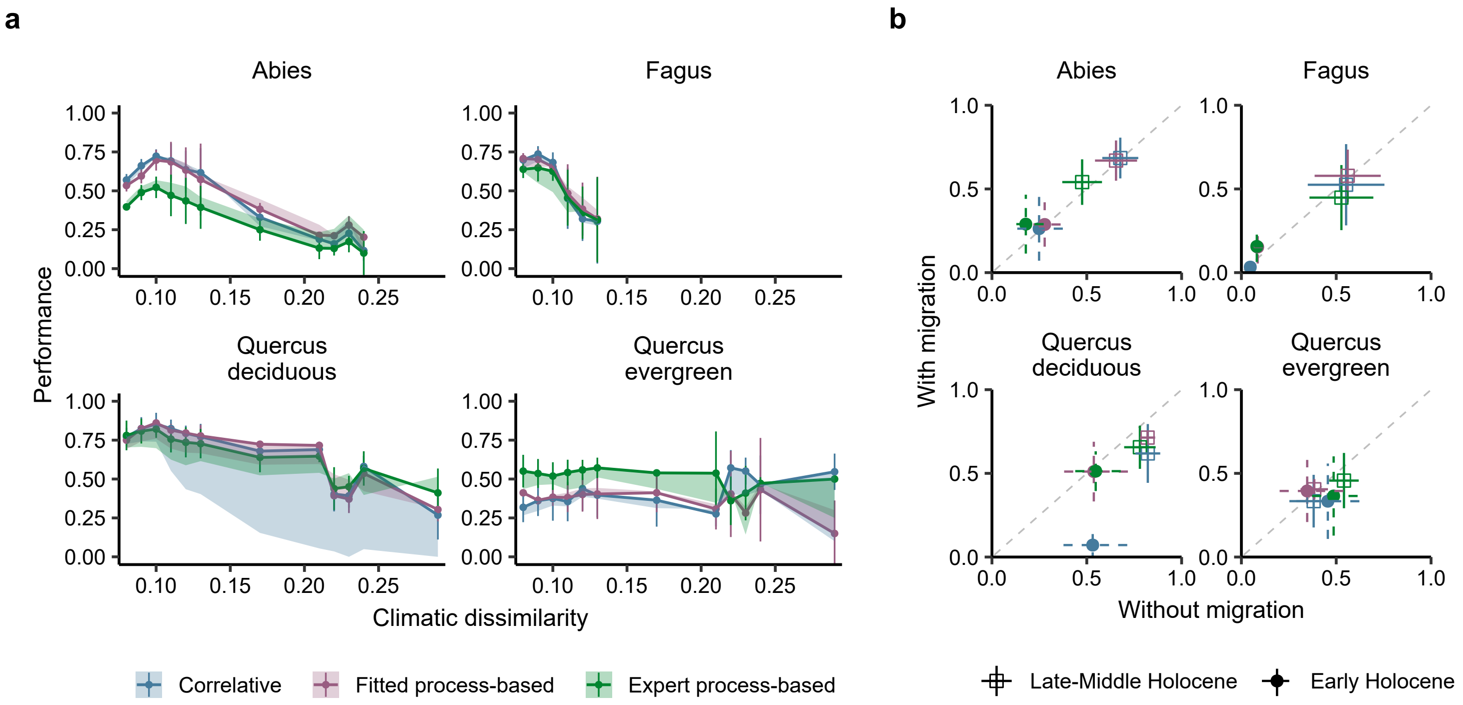


**Fig. 9 Impacts of migration on model performance, per species. (a)** Evolution of performance against climatic dissimilarity. Lines and points represent model performance without migration, shaded areas show performance evolution when simulating migration. Panel **(b)** displays model performance with and without migration, for Late-Middle and Early Holocene.

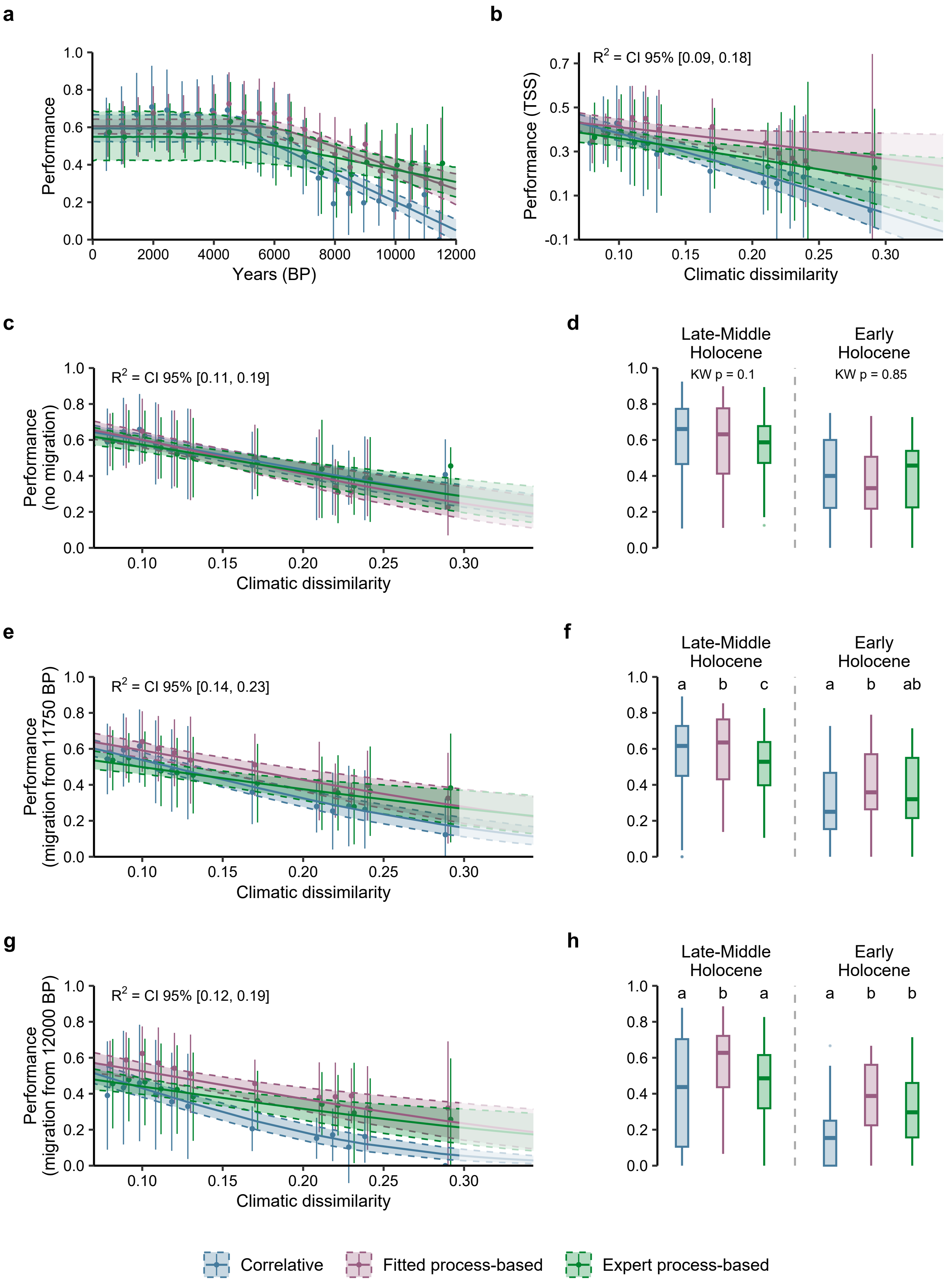


**Fig. 10 Performance of correlative models, fitted process-based models (inverse calibration using occurrence data) and expert process-based models (classical calibration). (a)** Model performances against time. Lines are linear-plateau regressions, which follow two phases (a flat plateau and a linear response). Shaded areas represents 2.5% and 97.5% confidence intervals, calculated with the R package *propagate* [5] by using first and second-order Taylor expansion and Monte Carlo simulations. **(b)** Bayesian beta regression of model performance (TSS) against climatic dissimilarity, with the same simulations as in main text (migration started from 12 kyr BP, or 11.75 kyr BP when a model simulates no presence at 12 kyr BP). **(c,e,g)** Bayesian beta regression of model performance (Sørensen index) against climatic dissimilarity, and **(d,f,h)** difference in performance across models. The grouping letters represent the multiple comparisons with pairwise Conover-Iman tests. "KW" stands for "Kruskal-Wallis". **(c,d)** Models without migration, **(e,f)** models with all migration starting points at 11750 BP, **(g,h)** models with all migration starting points at 12000 BP.

### References


[1] Andersen, K. K. *et al.* High-resolution record of Northern Hemisphere climate extending into the last interglacial period. *Nature* **431**, 147–151 (2004). URL https://www.nature.com/articles/nature02805.

[2] Armstrong, E., Hopcroft, P. O. & Valdes, P. J. A simulated Northern Hemisphere terrestrial climate dataset for the past 60,000 years. *Scientific Data* **6**, 265 (2019). URL https://www.nature.com/articles/s41597-019-0277-1.

[3] Burke, K. D. *et al.* Differing climatic mechanisms control transient and accumulated vegetation novelty in Europe and eastern North America. *Philosophical Transactions of the Royal Society B: Biological Sciences* **374**, 20190218 (2019). URL https://royalsocietypublishing.org/doi/10.1098/rstb.2019.0218.

[4] Thrasher, B. *et al.* NASA Global Daily Downscaled Projections, CMIP6. *Scientific Data* **9**, 262 (2022). URL https://www.nature.com/articles/s41597-022-01393-4.

[5] Spiess, A.-N. *propagate: Propagation of Uncertainty* (2018). URL https://CRAN.R-project.org/package=propagate. R package version 1.0-6.
